# Reciprocity and interaction effectiveness in generalised mutualisms among free-living species

**DOI:** 10.1101/2022.03.23.485462

**Authors:** Elena Quintero, Francisco Rodríguez-Sánchez, Pedro Jordano

## Abstract

Mutualistic interactions among free-living species generally involve weak links and highly asymmetric dependence among partners, yet our understanding of factors beyond their emergence is still limited. Using individual-based interactions of a super-generalist fleshy-fruited plant with its frugivore assemblage we estimate the Resource Provisioning Effectiveness (RPE) and Seed Dispersal Effectiveness (SDE) to assess the balance in the exchange of resources. Plants were highly dependent on a few super-generalist frugivore species, while these interacted with most individual plants, resulting in strong asymmetries in mutual dependence. Both RPE and SDE were mainly driven by interaction frequency. Despite highly asymmetric dependences, the strong reliance on quantity largely determined high reciprocity in rewards between partners (i.e., higher energy provided, more seedlings recruited), not obscured by minor variations in the quality of animal or plant service. We anticipate reciprocity will emerge in low-intimacy mutualisms where the mutualistic outcome largely relies upon interaction frequency.

## INTRODUCTION

Mutualisms are ecological interactions entailing beneficial outcomes for the interacting partners. These outcomes broadly emerge from the exchanges and “fair two-way transfer of resources” resulting from the interspecific encounters (Kiers et al. 2011). Despite recent interest in interspecific exchanges, especially focusing on strict and intimate interactions (Guimarães *et al*. 2007), much of the reciprocal effect between generalised, free-living, mutualistic partners remains unexplored (Thompson 2009).

Species-level analyses of complex interaction networks have revealed a highly heterogeneous structure (i.e., high variance in number of interactions per species), weak levels of mutual dependence, and high asymmetry in interaction strength (i.e., marked difference in partners’ dependencies for any pairwise interaction; Jordano 1987; Johnstone & Bshary 2008; Bascompte & Jordano 2014; Wootton & Stouffer 2016). Interaction asymmetry in complex networks of free-living species (Bascompte et al. 2006) as well as energy flow asymmetry in food webs (Rooney et al. 2006) appear as quintessential characteristics of these complex systems, closely associated with their stability. Yet our understanding of the factors beyond the emergence of asymmetric interactions is still very limited; for example, if any mutualistic interaction between free-living species entails an exchange of services, is there a “fair two-way transfer” of resources in these generalised mutualisms (Kiers et al. 2011; Chomicki et al. 2020), in other words, is there reciprocity?

Reciprocity, as defined herein, is the existence of a positive association in the rewards provided between mutualistic partners. We consider a mutualistic system to be reciprocal (to a varying degree) if higher reward provided by one organism (e.g., more pollen grains or fruits offered by plants) results in higher reward from its mutualistic partner (e.g., more fertilised ovules or dispersed seeds). In contrast, if higher rewards offered by one partner do not return increased rewards by the other partner (e.g., because offering more pollen or fruits attracts more antagonists, or mutualists cannot cope with the increased availability of resources), those interactions would be less, or not reciprocal at all. Without an external reference, reciprocity cannot be estimated directly, as it is not possible to determine if the exchange in resources between partners is equal or fair. Reciprocity can only be understood using a more general perspective by comparing the resource exchange between partners within a specific interaction relative to the general pool of interactions. A population or community perspective will allow us to understand whether specific pairwise interactions are exchanging their resources at ‘fair’ cost, or at least at the cost set by the population or community. Aside from previous work on mycorrhizal symbiosis, less intimate and ‘lagged’ (i.e., with delayed responses beyond the interaction itself) mutualisms that examine reciprocity as hereby defined have been rarely addressed. Notwithstanding, previous studies explore other concepts of reciprocity using different approximations, more related to the degree of partner’s dependence (see, e.g.: (Herrera 1984; Reid 1990; Burns 2003; Guerra & Pizo 2014).

A better explored aspect of mutualistic interactions is partner dependence, i.e., how much a partner relies upon another specific partner for their services. Dependence could be estimated as the proportion of service obtained from a specific partner relative to the total service obtained from the whole set of interactions. Dependence differs from reciprocity in that it examines the reliance from the perspective of the partner, and not the whole population. Estimating dependence also allows calculating the asymmetry of interactions, by comparing the mutual dependence of both partners in a mutualism. Thus, asymmetry shows up when a species/organism depends a lot on one partner but, in turn, this one does not rely too much on that particular pairwise interaction (Jordano 1987; Bascompte et al. 2006; Vázquez et al. 2007).

In fact a generalised property of free-living species networks is the high frequency of weak interactions (Jordano 1987) so that when interactions are strong, they are invariably highly asymmetric. This pattern in the mode of interaction between organisms is known as disassortativity, whereby organisms with high-degree tend to interact with organisms of lower-degree (Barabási 2016), and is recurrently found in biological networks (Newman 2003). Weak links appear a characteristic feature of complex systems made up of highly diversified components (Granovetter 1973; Csermely 2009) and provide support for their stability (McCann et al. 1998; Berlow 1999). Most previous analyses of network patterns in real-world ecosystems have considered species-level interactions. Yet, interaction asymmetries at the individual-level remain largely unexplored, despite likely being the most appropriate level to address interaction outcomes (Clark *et al*. 2011). Actual ecological interactions in nature that we can observe, sample, monitor, and document, occur from interspecific encounters among individuals of the partner species (Dupont et al. 2014; Jordano 2016). One might therefore wonder if, when looking at a more refined level (e.g., from species to individuals), we could still expect asymmetry in their mutual dependence.

Few studies so far have analysed interaction asymmetry beyond variation in just interaction frequency or strength, e.g., further examining differences in interaction quality (Herrera 1984; Jordano 1987; Guerra & Pizo 2014; González-Castro *et al*. 2022). Interaction outcomes may yield very different results from those expected solely on the basis of interaction frequency (Janzen 1983), and so it is possible that infrequent interactions result in higher fitness values than frequent interactions, affecting the reciprocity balance between the mutual dependencies. A useful tool to measure the functional outcome (fitness) of mutualistic relationships in terms of both interaction quantity and quality is the effectiveness framework (Schupp 1993; Schupp et al. 2017, Fig.1A). Considering individual variation and interaction outcomes expands our understanding of the potential consequences, e.g., demographic or evolutionary, that depend on fitness variation among individual partners, especially when effectiveness and its components are estimated for both the plant and animal species sets.

**Figure 1.**
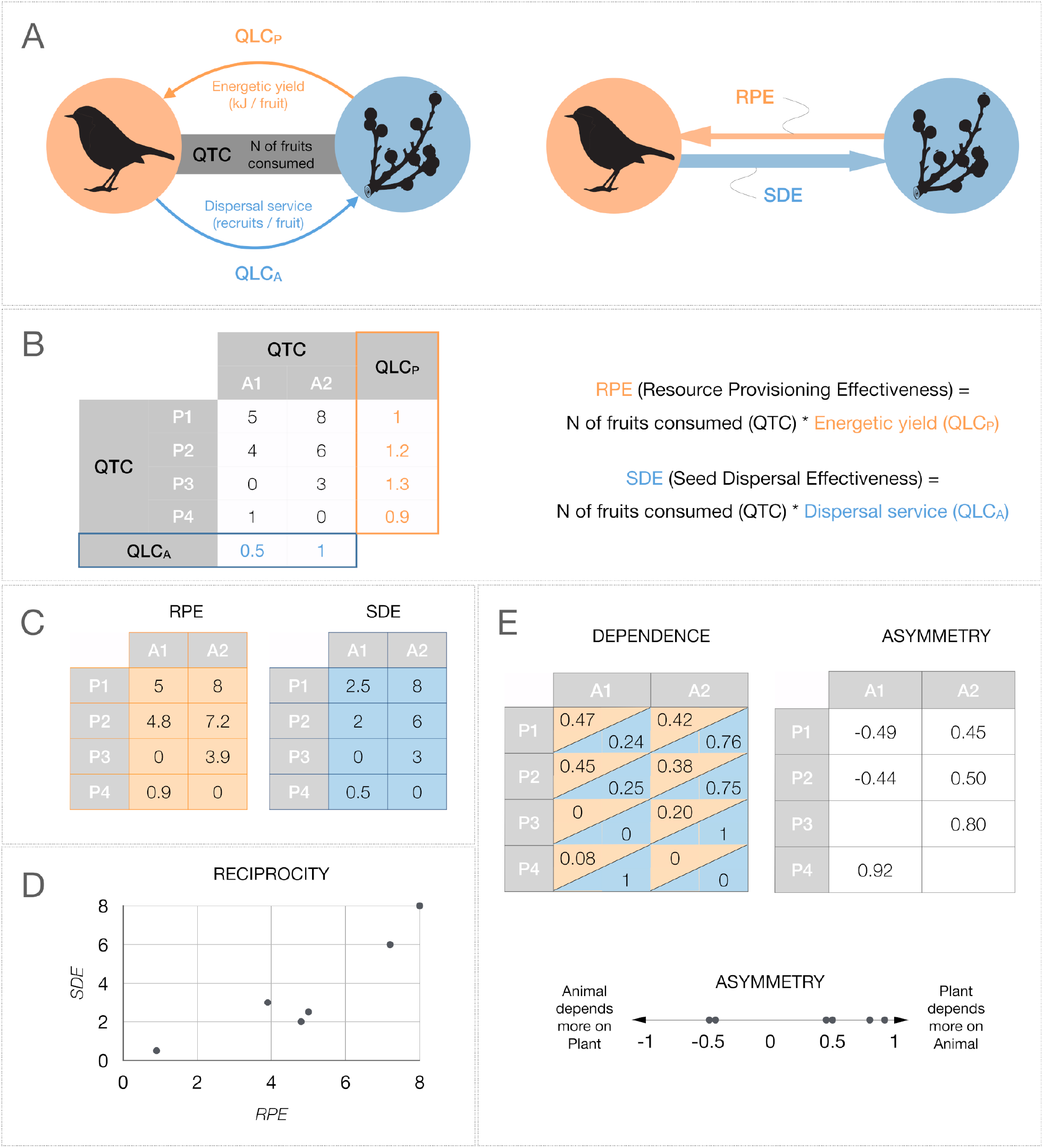
Schematic representation of the approximation used in this study for the characterization of plant-frugivore seed dispersal mutualisms. (A) the three main subcomponents which are present in the mutualism between any two nodes (animal frugivore, orange; plant individual, blue) in the network: the interaction frequency or quantity (QTC) and the two-sided quality (QLC) of the service provided by the partner: energetic yield provided by the plant (QLC_P_) and probability of seedlings recruitment provided by the bird (QLC_A_). The effectiveness for the two partners (seed dispersal for the plant: SDE, and resource provisioning for the animal: RPE) is estimated by combining these subcomponents. (B) An example adjacency matrix of quantity and quality data for the calculation of effectiveness (left) and the expressions to derive RPE and SDE (right). (C) Resulting calculations of RPE and SDE using the example matrix in B. (D) Reciprocity can be assessed by the covariation between RPE and SDE values and characterises the sign and direction of the outcome of any pairwise interaction. (E) Derivation of mutual dependence estimates and interaction asymmetry obtained for plant and animal partners. Dependence values for animals in the orange upper-left cell are calculated based on RPE values, while dependence values for plants in the blue lower-right cells are calculated based on SDE values (left matrix, see eq. 1 below for calculations). With both estimates of the mutual dependence matrix it is possible to calculate their asymmetry (right matrix, see eq. 2 below for calculations). Asymmetry is estimated as a standardised difference between the two dependence values in each interaction, and ranges between -1 and +1.

In this study we calculate the two-sided rewards for seed dispersal mutualistic interactions between plants and animal frugivores by means of the Resource Provisioning Effectiveness (RPE) and Seed Dispersal Effectiveness (SDE) frameworks. We look at mutual reciprocity (i.e., the balance in the exchange of resources) from an individual perspective in a plant population using SDE and RPE as estimates for the reward obtained in the relationship (Fig. 1D). We also explore if mutualistic dependencies are still asymmetrical when looking at a plant individual perspective and when incorporating not only the frequency of the interactions, but also their quality (i.e., interaction outcome) (Fig. 1E). For this purpose we use as study organism the super-generalist plant species *Pistacia lentiscus* (Anacardiaceae). Super-generalist species interact with a large part of the local diversity of partner species and connect semi-independent modules in the community, conferring them a fundamental role in ecological networks as they provide great cohesion (Guimarães et al. 2011). The analysis of individual-based, pairwise interactions thus allows a direct link to evolutionary approaches based on empirical data of fitness variation in relation to phenotypic traits and the interactions modes of individuals as a basis to understand natural selection in mutualisms. A two-sided study of mutualism at this individual level provides us with information on the diversity of individual plant rewards, the diversity of mutualistic partners and their effects, and the consequences on resource exchange between them.

Here we address these specific objectives: 1) characterise the effectiveness of the mutual beneficial service between individual plants and their frugivorous species, 2) test if the service provided between partners in terms of the amount of reward is reciprocal, and 3) explore if there exists asymmetry in the mutual dependencies when looking at a plant individual level and considering interaction outcomes, that is, accounting for interaction quality beyond interaction frequency.

## METHODS

### Species and study site

*Pistacia lentiscus* (Anacardiaceae) is a dioecious and anemophilous pollinated shrub that can be considered as a ‘foundation species’ (Whitham *et al*. 2006) playing a central role in the landscape physiognomy of lowland Mediterranean scrublands. Numerous resident and migrant frugivorous birds rely on lentisc fruits as a nutritional resource (González-Varo *et al*. 2019) and act as its seed dispersers, with infrequent consumption by mammals (Perea *et al*. 2013).

Fieldwork was conducted at two study sites in Doñana National Park (Huelva, SW Spain): La Mancha del Rabicano in El Puntal site (EP) and Laguna de las Madroñas (LM). Both areas consist of Mediterranean sclerophyllous scrubland dominated by lentiscs (*Pistacia lentiscus)* coexisting with a total of 28 fleshy-fruited species recorded in the area.

A total of 80 individual lentisc plants were marked, 40 per study site (Suppl. Mat. A). In order to estimate the resource provisioning and seed dispersal effectiveness (RPE and SDE, Fig. 1), we studied the frequency (i.e., the quantity; QTC) and the functional outcome (i.e., the quality; QLC) of the pairwise interactions between the individual lentiscs and the avian frugivores present in the area.

### Interaction frequency: QTC

The interaction frequency of *Pistacia lentiscus* plants with avian frugivore species was assessed through DNA-barcoding techniques and continuous-monitoring cameras (Quintero *et al*. 2021; Suppl. Mat. B). Individual plant monitoring took place during the complete fruiting season, between September 2019 and March 2020.

For molecular DNA analysis, we placed seed traps beneath individual plants, where we collected a total of 2691 faecal and seed samples (1913 for EP and 778 for LM). Identification of visiting species was done applying DNA-barcoding analysis to collected samples. Animal-origin DNA present in the surface of the samples was extracted, amplified and then sequenced following protocol in González-Varo *et al*. 2014 with minor modifications (Suppl. Mat. B.1). Retrieved sequences were identified using the BOLD Systems database (https://www.boldsystems.org/) or the BLAST from the NCBI (https://blast.ncbi.nlm.nih.gov/Blast.cgi). Identification success rate of the analysed samples was 94% (n = 2285).

With monitoring cameras we recorded animal visitation and feeding events in focal plants at EP site. All individual plants were monitored every fortnight along the fruiting season, accumulating c.19 h observation per plant (Suppl. Mat. B.2). Video recordings were analysed with the help of the motion detection program DeepMeerkat (Weinstein 2018; Suppl. Mat. B.2). We obtained the feeding frequency of animal species (i.e. fraction of visits with actual fruit consumption) and the number of fruits consumed per visit. Overall, cameras recorded 3790 visits. Species identification was possible for 91% of the visits (n = 323 visits by unknown species). A total of 37 animal species were identified to be interacting with the individual plants of which 26 species were frugivorous birds and 24% of them included apparent feeding records.

The total number of frugivorous species recorded was 27; 26 recorded with cameras and 22 with DNA-barcoding. Interaction accumulation curves (IAC) were used to determine both DNA-barcoding and video recording sampling completeness (see Suppl. Mat. B.3; Colwell & Coddington 1994; Jordano 2016). Overall sampling completeness was 93% for both methods (sensu Chacoff *et al*. 2012); 95% just for cameras and 96% just for DNA-barcoding.

To estimate the total number of fruits consumed by each bird species at each individual plant (Fig. 2) we multiplied these four sequential steps: (1) the total number of visits at each site, (2) the probability that a given bird species visits a particular plant, (3) the probability that a visit includes a feeding event, and (4) the number of fruits/seeds consumed per visit by each bird species. We estimated these quantities using Bayesian models fitted with Stan (Stan Development Team 2022) and brms (Bürkner 2017) (Suppl. Mat. E.1). This approach allowed us to account for more realistic estimates of uncertainty and obtain probabilistic estimates for few unobserved quantities.

**Figure 2.**
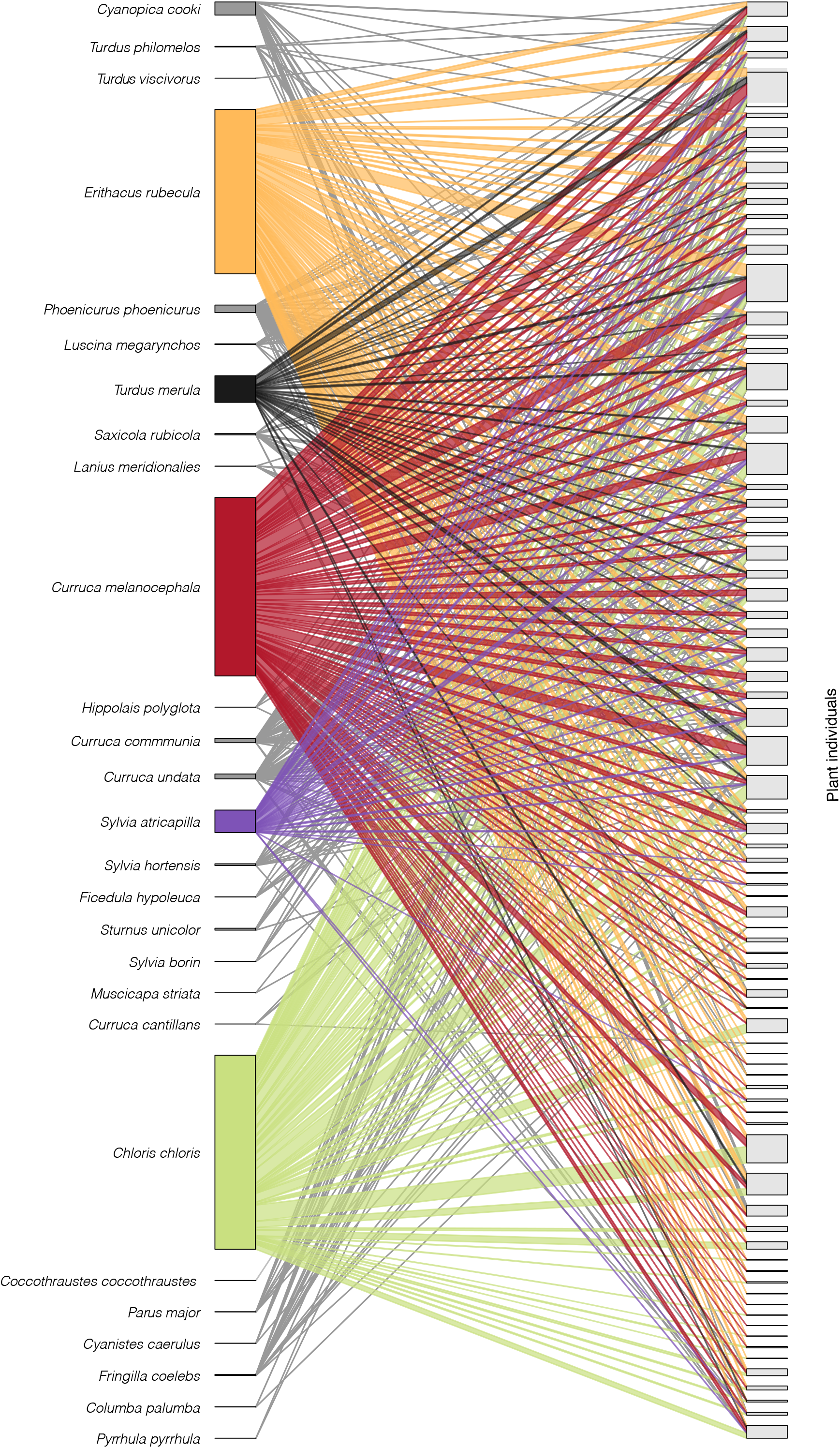
Representation of interaction network between avian consumers species and individual *Pistacia lentiscus* plants, where the node and link width is proportional to the total number of fruits consumed on each plant. Non-legitimate dispersers (n=7) are grouped at the left end of the network.

### Interaction outcome for the animal: QLC - RPE

Plant quality was defined as the energetic reward provided per fruit or seed (for granivorous birds; see Table S.A.1 for frugivory type categories). We collected fruits from each plant (mean = 31 fruits, range = 17-63, Suppl. Mat. C) and measured both pulp and seed fresh mass. Fresh mass was converted to dry mass using *P. lentiscus* water % content (Jordano 1984). We then multiplied the pulp and seed dry mass by their estimated energy yields: 25.25 kJ/dry g of pulp and 28.14 kJ/dry g of seed (see Suppl. Mat. E.2; Herrera 1987; MacLean *et al*. 2003; Khiari *et al*. 2020).

### Interaction outcome for the plant: QLC - SDE

We used Bayesian models (Suppl. Mat. E.3) to estimate the quality of animals as seed dispersers according to: (1) probability of seeds to escape granivorous bird predation during handling, (2) microhabitat use by each bird species, (3) probability of seeds escaping rodent post-dispersal predation, and (4) probability of seedling emergence and early survival (past their first summer) in each specific microhabitat. The product of these steps rendered the probability of seedling recruitment resulting from the consumption of one fruit by a specific avian consumer.

The probability of seeds to escape bird predation was estimated by counting the number of intact seeds manipulated by predators (identified through DNA-barcoding) collected in seed traps beneath plants (Suppl. Mat. D.1). Microhabitat use by different bird species was inferred from the seed rain of *P. lentiscus* seeds collected at five microhabitats: under *Pistacia lentiscus* conspecifics (PL), under other fleshy fruited species (FR), under non-fleshy fruited species (NF), under pine trees (*Pinus pinea*; PP), and open ground areas (OA) (see Suppl. Mat. D.2). We identified the bird species through DNA-barcoding of collected seeds. For each microhabitat we also measured post-dispersal predation, seedling emergence and survival through seed removal and seed-sowing experiments (Suppl. Mat. D.3, D.4).

### Effectiveness calculations

We calculated the final effectiveness as the product of quantity and quality components (Suppl. Mat. E; Fig. S.E.1). The quantity component (i.e., total number of fruits consumed by a specific bird on a given plant) was common for both the animal and plant’s perspective. Quality for the animal was the energy acquired per fruit/seed consumed. Quality for the plant was the probability that a consumed fruit becomes a seedling surviving its first summer. Resource Provisioning Effectiveness (RPE) therefore estimates the total energy provided by each plant to each bird species along the fruiting season, and Seed Dispersal Effectiveness estimates the potential number of seedlings recruited by each plant through interacting with each bird species.

### Reciprocity

To estimate the reciprocity we used Pearson correlation coefficients between the log-transformed RPE and SDE values. We aggregated the total rewards offered and received by each individual plant across all bird species, using the 1000 posterior distribution samples (see Suppl. Mat F.1). A high positive correlation would indicate high reciprocity, meaning that plants providing high resource provisioning (RPE) obtain in turn high dispersal effectiveness (SDE).

### Calculating dependence and asymmetry between individual plants and bird species

We calculated mutual dependence (*d*) for each pairwise interaction (Suppl. Mat. F.2). Two separate dependence values were obtained, one for the plant 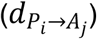 and one for the animal species 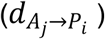.

eq. 1a: 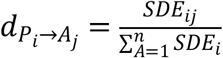, for the dependence of *P. lentiscus* plant *i* on animal species *j*; and eq. 1b: 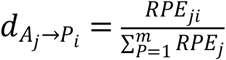, for the dependence of animal species *j* on plant *i*, where *d* is the dependence of plant *i* on animal species *j*, or vice versa; *SDE*_*ij*_ is the estimated number of seedlings recruited by plant *i* via frugivore species *j*; *RPE*_*ji*_ is the amount of kilojoules plant *i* provided to frugivore species *j*; and *n* and *m* represent the total number of animal species and individual plants, respectively.

Interaction asymmetry (AS) is defined as (Bascompte *et al*. 2006; Vázquez *et al*. 2007):

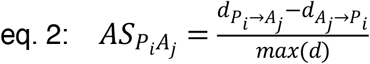

AS values can range from -1 to 1, where 0 indicates total symmetry (i.e. both partners depend on each other with the same intensity), values approaching +1 indicate that the plant is more dependent on the animal than *vice versa*, and negative values indicate that the animal is more dependent on the plant than the plant on the animal.

We performed different null models to test the robustness of the observed asymmetry values to our sampling design (see Suppl. Mat. H). We did not find evidence for asymmetry values being significantly biased in any of these null models.

## RESULTS

### Plant individual-based interactions

We estimated that birds consumed a total of 2.2 × 10^5^ fruits from the 80 marked plants at both *P. lentiscus* populations (90% credibility interval: 1.5 × 10^5^ - 6.6 × 10^5^). This represents ∼20% of the total number of fruits produced by these plants in the 2019-20 season (Supp Mat G.1). We detected 27 bird species consuming *P. lentiscus* fruits, of which 12 are considered residents, 9 summer or trans-Saharan migrants and 6 winter migrants (see Suppl. Mat. A). More than 85% of the consumed fruits were consumed by just three species, *Curruca melanocephala, Erithacus rubecula* and the seed predator *Chloris chloris*. These species behaved as super-generalists, interacting with the great majority of individual *P. lentiscus* plants (see Fig. 2). The next stronger consumers were *Turdus merula* and the winter migrant *Sylvia atricapilla*.

### Resource Provisioning and Seed Dispersal Effectiveness

*Pistacia lentiscus* plants were highly variable in the Resource Provisioning Effectiveness (RPE) provided to avian species (Fig. 3). Bird species consumed a median of 97 fruits/seeds on each plant (interquartile range: 23 - 474). We estimated that *Curruca melanocephala* and *Erithacus rubecula* may have eaten more than 4000 fruits, and *Chloris chloris* more than 5500 seeds, at some individual plants. This intensity of consumption represents, however, just a small proportion of the available crop offered: most plants had less than half their crop size removed by birds (see removal success in Suppl. Mat. G.1). The quantity component accounted for almost all (93%) of the variation in RPE (Suppl. Mat. S.E.5). Regarding quality, we found up to 7-fold differences in the energetic content of fruits/seeds from individual plants. For fruit/pulp consumers, quality ranged from 0.11 to 0.77 kJ/fruit, whereas seed predators obtained between 0.11 and 0.66 kJ/seed. Birds consumed fruits and seeds of varied quality within those ranges, following energy availability (Suppl. Mat. G.2). In general, avian consumption was higher in plants with larger crops, canopy area, and pulp content (Suppl. Mat. G.3).

**Figure 3.**
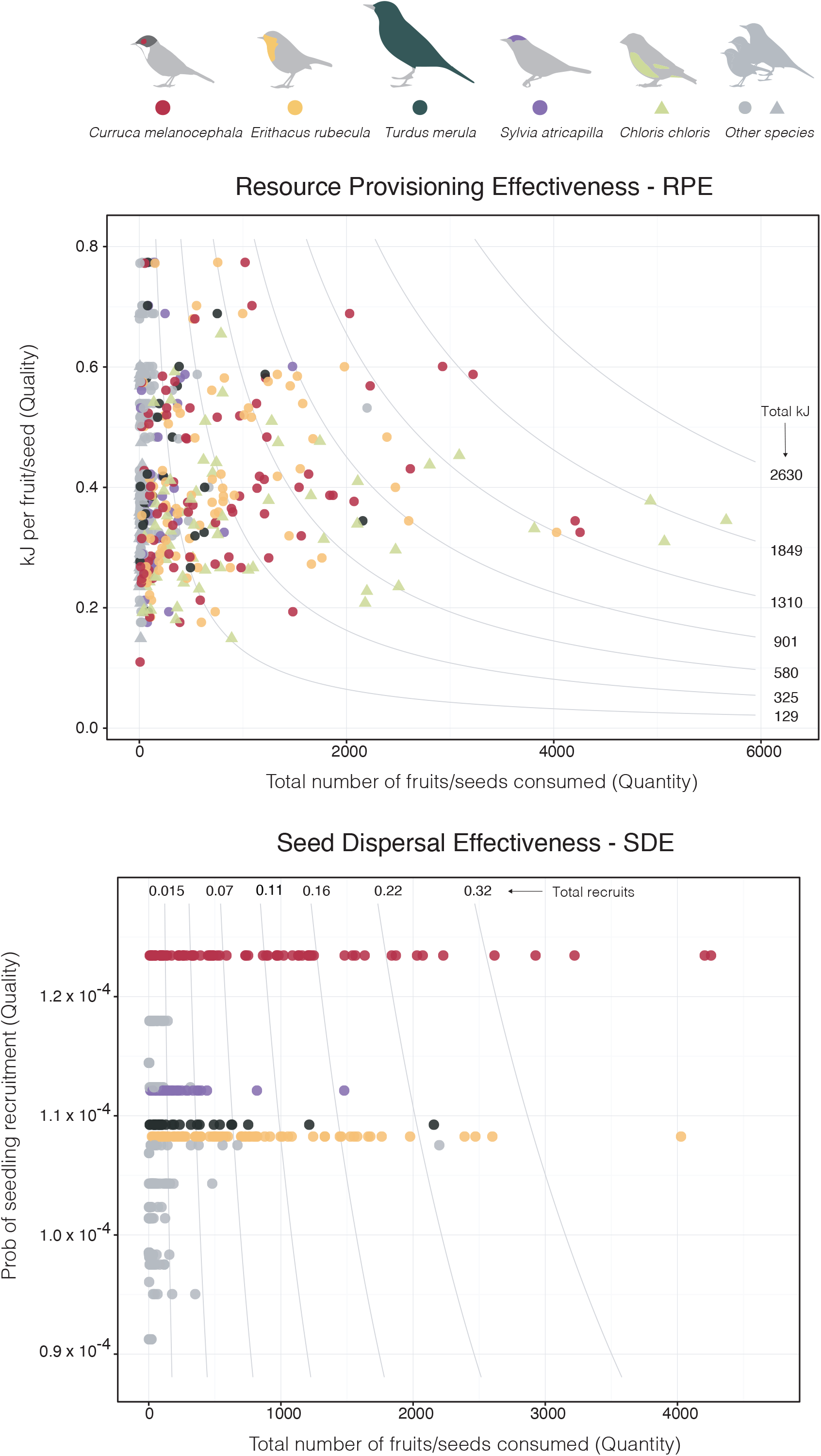
Effectiveness landscapes for resource provisioning (RPE) and seed dispersal (SDE). Each point represents an individual pairwise interaction. In both landscapes, the horizontal axis depicts the total number of fruits/seeds consumed by each bird species in each individual plant. In the RPE landscape, the vertical axis represents the median energy (kJ) obtained from a fruit/seed from each individual plant. In the SDE plot, the vertical axis represents the posterior median probability of recruiting a seedling from a fruit ingested by each bird species. Hence the product of the horizontal (Quantity) and vertical (Quality) axis gives the effectiveness of each bird-plant pairwise interaction: the total energy (kJ) in the case of RPE, and total number of plant recruits for SDE. Different combinations of quantity and quality can produce equal effectiveness values, as shown by isolines. Note seed predators are not shown in the SDE landscape visualisation, as their dispersal quality is zero or close to zero and their inclusion distorts the graph (see Supp Mat E.4 for complete SDE landscape).

Seed Dispersal Effectiveness (SDE) was also more determined by the quantity (fruit/seed consumption) than the quality component (probability of seedling recruitment), which varied little among bird species (variance partitioning: quantity = 69%, quality = 31%; Suppl. Mat. E.5) (Fig. 3). Excluding seed predators, with negligible contributions to recruitment (as they destroyed ∼99.9% of the seeds consumed), all bird species had a similar probability of producing a seedling surviving the first summer drought (around 10^−4^ per consumed fruit), with *Curruca melanocephala* emerging as the highest quality disperser, followed by other members of the Sylviidae family. Differences among frugivore species in dispersal quality stem from their distinctive microhabitat use (Suppl. Mat. E.3.1) and existing trade-offs between recruitment stages in different microhabitats (Suppl. Mat. E.3.2; E.3.3). For example, seeds falling under *Pinus pinea* trees had the highest probability of surviving rodent predation, followed by those arriving to open areas. Seedling emergence and survival, on the other hand, was highest in open areas and lowest beneath *Pinus pinea*. Overall, open area was the microhabitat with highest probability of recruitment, yet very few seeds arrived to it, hence this microhabitat hardly contributed to recruitment. The relatively high quality of *C. melanocephala* emerged from its preferential dispersal towards the most suitable microhabitats: beneath non-fleshy fruited plants and *P. pinea*. In contrast, heavy *P. lentiscus* fruit consumers like *E. rubecula* showed medium quality as it frequently deposits seeds under *P. lentiscus* plants, a microhabitat where the probability of escaping post-dispersal seed predation and seedling survival were medium-low.

Despite the quite high fruit consumption, overall probabilities of recruitment at the final stage considered in our study (i.e., seedlings surviving the first summer drought) were rather low. Even the most intense pairwise interaction observed (involving *C. melanocephala*) would have recruited roughly half a seedling surviving its first summer (SDE value of 0.53).

### Reciprocity

We found high reciprocity in the interactions between individual *P. lentiscus* plants and their bird consumers: on average, plants supplying more energy (i.e., having more fruits/seeds consumed) also recruited a larger number of seedlings (Fig. 4). This is supported by the high correlation between RPE and SDE (mean Pearson r on log-log values = 0.93; mean 90% CI = 0.90 - 0.96; see Suppl. Mat. F.1). In other words, the larger the reward provided by one interaction partner (the plant), the larger the reward contributed by the other partner (birds). This high reciprocity stems from the fact that both RPE and SDE were mainly driven by the quantity component (i.e., intensity of consumption) rather than by differences in plant and frugivores quality. As a result, plants mobilising more fruits also recruited more seedlings (on average), regardless of differences in the composition of their frugivore assemblages. Additionally, plants involved in greater rewards tended to have larger crop sizes and were consumed by a higher number of bird species.

**Figure 4.**
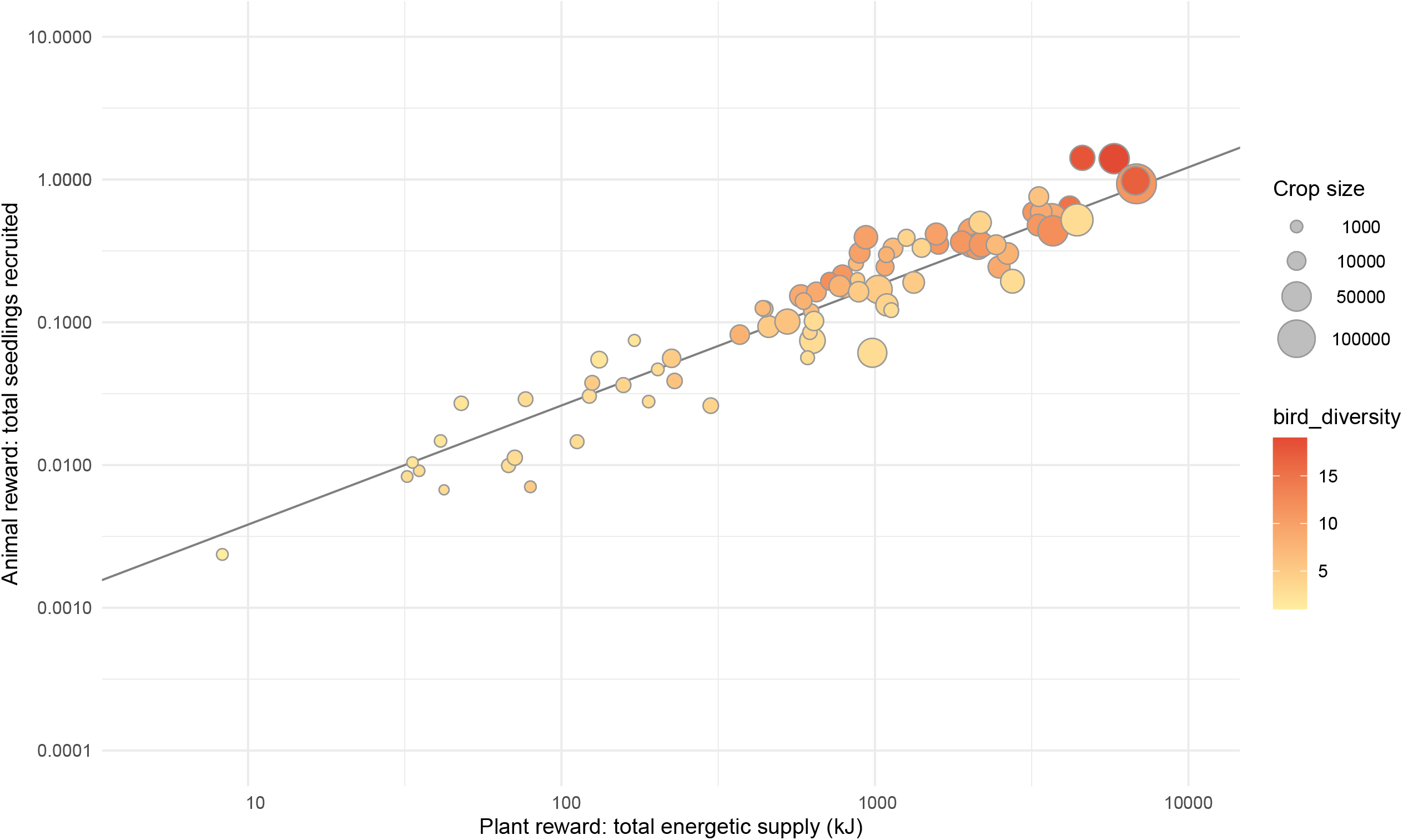
Relationship between the total energetic supply provided by individual plants (aggregating all its consumer bird species) and the number of seedlings recruited by each plant (n = 79). The positive relationship indicates highly reciprocal interactions: the higher the reward offered by the plant (i.e., more fruits consumed), the higher the reward received from its bird consumers. Dot size represents plants’ initial crop size and colour intensity indicates the number of animal species partners. Note both axes are in logarithmic scale.

### Dependence and Asymmetry

Dependencies on the mutualistic partner were in general low (Fig. 5). Most bird-plant pairwise interactions had dependencies below 0.25, meaning that most interactions actually reported only a small fraction of the total reward (i.e., energy income or seedlings recruited) for either partner (birds and plants, respectively). There were, however, some strong, highly-dependent interactions, namely those involving the two main dispersers *E. rubecula* and *C. melanocephala*: most plants strongly depended on both bird species for effectively dispersing their seeds and recruiting (Fig. 5, left). In contrast, avian species were remarkably less dependent on individual plants. Only a few rare bird species (e.g. *Turdus viscivorus* and *Hippolais polyglotta* among fruit consumers, and *Coccothraustes coccothraustes* and *Pyrrhula pyrrhula* among seed predators) showed high dependency on specific plants (Fig. 5, centre).

**Figure 5.**
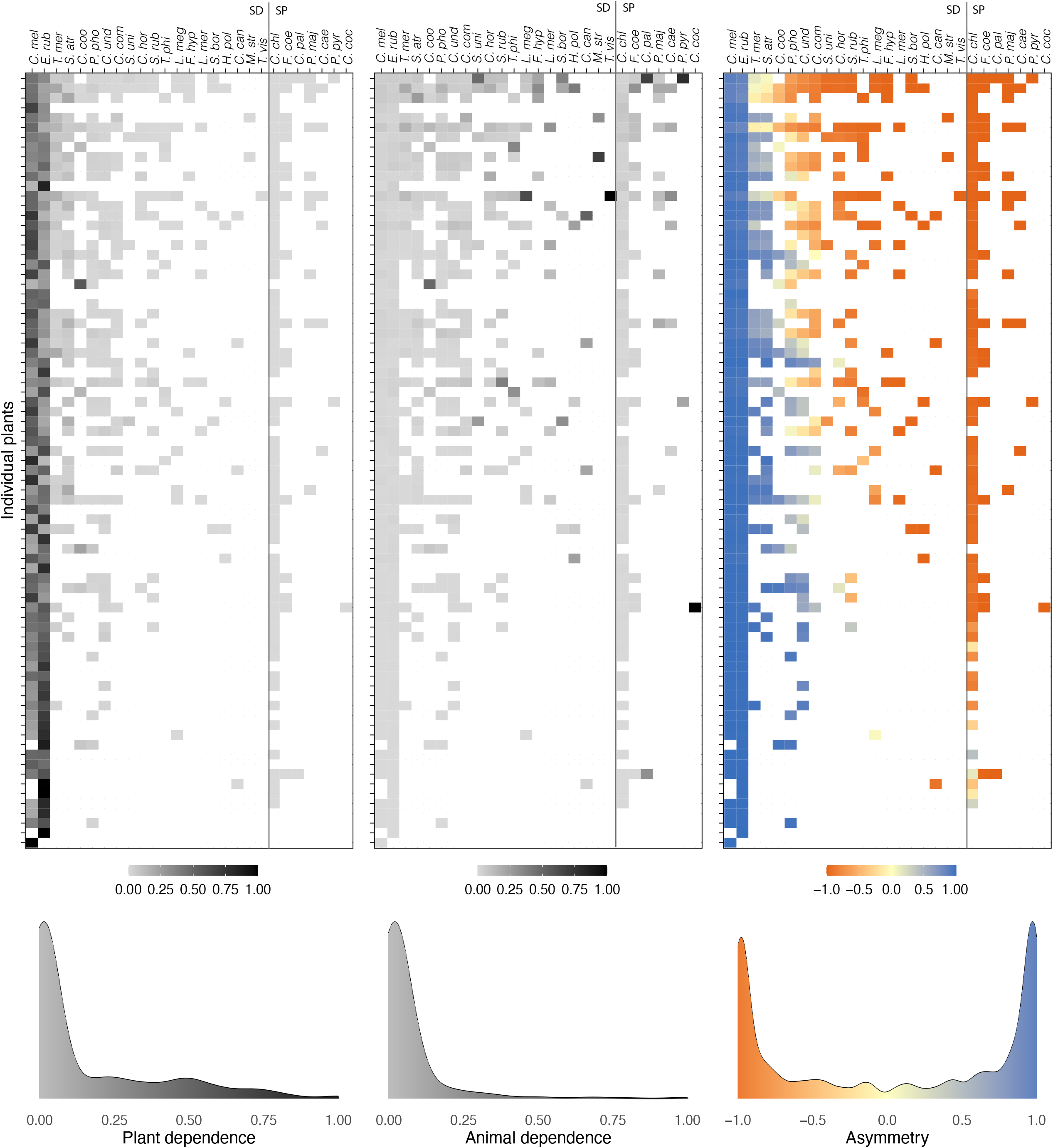
Interaction matrices between *Pistacia lentiscus* individual plants and their avian consumers. The first matrix (left) depicts how much each plant seed dispersal effectiveness (number of seedlings recruited) depends on each bird species, whereas the second matrix (center) shows how much the resource provisioning effectiveness (energy obtained) of each bird species depends on each particular plant. Both matrices range from 0 (no dependence at all) to 1 (total dependence on that particular partner). The third matrix (right) shows the asymmetry in dependence for each unique bird-plant pairwise interaction. Colours gradually veering toward blue (asymmetry values approaching 1) indicate interactions where the plant is more dependent on the animal than vice versa, whereas colours veering toward orange (i.e., asymmetry approaching -1) indicate interactions where the animal is more dependent on the plant. Symmetrical interactions, where the dependence of both partners is similar, are represented by yellow tones (asymmetry values close to 0). The lower graphs represent the frequency distribution of the above matrix values. Animal species codes in alphabetical order: *C*.*cae = Cyanistes caeruleus, C*.*can = Curruca cantillans, C*.*chl = Chloris chloris, C*.*coc = Coccothraustes coccothraustes, C*.*com = Curruca communis, C*.*coo = Cyanopica cooki, C*.*hor = Curruca hortensis, C*.*mel = Curruca melanocephala, C*.*pal = Columba palumbus, C*.*und = Curruca undata, E*.*rub = Erithacus rubecula, F*.*coe = Fringilla coelebs, F*.*hyp = Ficedula hypoleuca, H*.*pol = Hippolais polyglotta, L*.*meg = Luscinia megarhynchos, L*.*mer = Lanius meridionalis, M*.*str = Muscicapa striata, Pmaj = Parus major, P*.*pho = Phoenicurus phoenicurus, P*.*pyr = Pyrrhula pyrrhula, S*.*atr = Sylvia atricapilla, S*.*bor = Sylvia borin, S*.*rub = Saxicola rubicola, S*.*uni = Sturnus unicolor, T*.*mer = Turdus merula, T*.*phi = Turdus philomelos, T*.*vis = Turdus viscivorus*.

When comparing the corresponding dependencies of each partner, we found that most bird-plant interactions were highly asymmetric (Fig. 5, right) for two main reasons. First, most plants depended strongly on the main avian consumers (*C. melanocephala, E. rubecula*) while these birds had low dependencies on each individual plant (asymmetry values towards 1). That is, most individual plants required the service of these two frugivores for effective dispersal and recruitment, whereas these birds were feeding and obtaining energy from many plants, thus hardly depending on any particular one. Second, when the animals had high dependency on a particular plant (asymmetry values towards -1), the plants in turn hardly depended on that particular bird. This is explained by the fact that such interactions were dominated by seed predators, pulp consumers, and locally uncommon bird species, which provided no or very low seedling recruitment for plants. Symmetric interactions (where both partners had similar dependency values) were scarce and represented by strongly frugivorous and moderately abundant birds such as *T. merula, S. atricapilla* and *Cyanopica cooki*. In these cases, individual plants were similarly important for energy provisioning for these birds as they were for providing effective dispersal to plants.

## DISCUSSION

We report interaction patterns for a super-generalist plant species, with the aim to document variation in mutual dependence with animal seed dispersers at the plant individual level and degree of interaction reciprocity at the population scale. This allowed us to establish a proper link between the structure of individual-based interaction networks and the consequences in terms of fitness and local plant population recruitment.

### Interaction intensity dominates partner effectiveness

Most previous studies have dealt with interaction effectiveness from a species-level, community perspective (but see Guerra *et al*. 2017; Palacio 2019; Jácome-Flores *et al*. 2020). The individual focus in *P. lentiscus* has revealed ample variation in fruit consumption by animal frugivores at individual plants, while documenting smaller variances in the quality of partner’s reward.

Consumption intensity disproportionately affected the magnitude of the partner’s effectiveness. Both RPE and SDE variation were driven by the quantity component, rather than quality, with a 7-fold difference in individual energy content and a ∼1.3-fold variation in frugivore quality, but then three orders of magnitude variation in fruit consumption (quantity). This indicates that interaction frequency per se is acting as a good surrogate of effectiveness, as found in previous studies (Vázquez *et al*. 2005). Yet accounting for interaction quality may change interpretations of partner effectiveness in other systems (e.g. rank reversals in González-Castro *et al*. 2022).

The resource provisioning effectiveness landscape (RPE, Fig. 3) did not reflect clear preferences of bird species for plants with energy-rich fruits. However, when aggregating the consumption of non-granivorous birds by individual plants, we found that large plants, with larger fruit crops, producing heavier (more energetic) fruits, dispersed a larger number of fruits overall (Suppl. Mat. G.3). These traits are well known to affect frugivory (Sallabanks 1993; Ortiz-Pulido *et al*. 2007; Schupp *et al*. 2019) and are as well related to the ontogeny, growth and size hierarchies in plant populations (Weiner & Solbrig 1984). Other factors not analysed here, such as secondary compounds, fruit accessibility or fruiting neighbourhood could also be affecting consumption patterns (Moermond & Denslow 1985; Cipollini & Levey 1997; Carlo *et al*. 2007; Tonos *et al*. 2021).

Legitimate seed dispersers also exhibited limited variation in the quality component of seed dispersal effectiveness (SDE, Fig. 3). The resulting probability of recruitment was surprisingly similar between frugivore species, indicating a broad functional redundancy in their dispersal service (González-Castro *et al*. 2015). However, when considering the final effectiveness, two bird species (*C. melanocephala* and *E. rubecula*) emerged as the main contributors to seedling recruitment due to their high consumption. The redundancy encountered in the quality component would make the dispersal mutualism more robust to the loss of bird species, or fluctuations in bird populations (see Zamora 2000); yet marked changes in bird abundance, particularly of the dispersers that consume the most, could compromise plant recruitment.

### Reciprocity in partner rewards as a feature of mutualistic systems

Although the exchange of rewards between bird species and individual plants varied over several orders of magnitude, there was a high correlation between the rewards obtained by each partner in the interaction. This result points to a stable and fair two-way transfer in the exchange of mutualistic services. In the case of *P. lentiscus*, the reciprocity in the rewards stems from the strong dominance of the quantity (i.e., intensity of consumption), a common component on both resource provisioning and seed dispersal effectiveness. Such high reciprocity would appear characteristic of many seed dispersal systems and other generalised, resource-based mutualisms (Wheelwright & Orians 1982; Ollerton 2006). Yet, reciprocity in a mutualistic system could be compromised whenever there are large differences between partners quality (i.e., fruit energetic content, or recruitment probabilities for different dispersers), as occurs for example in systems with highly heterogeneous frugivore assemblages (González-Castro *et al*. 2015; García-Rodríguez *et al*. 2021). Reciprocity can also break down when antagonists differently disrupt mutualistic interactions of plants with legitimate seed dispersers (Jácome-Flores *et al*. 2020); yet mutualism breakdown scenarios have been largely examined for intimate interactions, not for free-living species (Sachs & Simms 2006; Chomicki & Renner 2017).

Aside from the high overall reciprocity, we found a ‘diminishing return’ effect, so that the number of seedlings recruited did not increase in the same proportion as the total energy provided by plants (mean slope of log SDE vs log RPE and SD = 0.83 ± 0.06; Fig. 4). This diminishing return in the number of seedlings recruited per unit of energy was not caused by interactions with seed predators (slope of the log-log relationship excluding seed predators: 0.85). *Chloris chloris*, the most frequent seed predator, attacked all plants in similar proportion. Instead, the deviation from strict proportionality (log slope = 1) could be caused by (i) plants producing heavier fruits disperse fewer seeds and recruit fewer seedlings per amount of energy offered than small-fruited plants, (ii) highly fecund individuals (dispersing many fruits) attracting both highly effective and less effective frugivores, and (iii) the fact that our analysis did not account for likely increasing recruitment probabilities with increasing fruit and seed size. If more energetic fruits containing more pulp also imply larger seeds with higher survival probability after dispersal (Piper 1986; Leishman *et al*. 2000), then our analysis would be underestimating the number of seedlings recruited for those plants. Our results are consistent with previous reports evidencing that extremely high seed production and consumption are required to ensure recruitment by mother plants, given the sharp decreases in survival probability as seeds move along dissemination and establishment stages (Herrera *et al*. 1994; García-Fayos & Verdú 1998; Gómez-Aparicio 2008). Following our estimates, individual lentisc plants would have to disperse >8000 seeds to have just a single recruit surviving their first summer. Thus, ensuring plant recruitment may require huge reproductive effort from plants, even in well-functioning dispersal mutualisms with high reciprocity.

### Highly asymmetric dependencies between mutualistic partners

The vast majority of interactions between bird species and *P. lentiscus* individual plants were highly asymmetric: when one partner depended strongly on some partner, the latter hardly depended on the former. The dominance of asymmetric interactions was driven, first, by the strong dependence of nearly all *P. lentiscus* plants on the most effective dispersers (*C. melanocephala* and *E. rubecula*) whereas these birds consumed fruits profusely from all plants, hence hardly depending on any particular one. Second, seed predators (remarkably *Chloris chloris*) depended on *P. lentiscus* plants for their energy income but did not in turn contribute new seedlings, generating another set of asymmetric interactions. Finally, some occasional fruit consumers focused on a few plants which hardly depended on these birds for recruitment. The highly skewed distribution of dependence values was likely generated by the combination of varying bird abundances (Vázquez *et al*. 2007) and degree of frugivory, plus varying fruit production and attractiveness to seed predators and legitimate seed dispersers from the plant side.

The high asymmetry between mutualistic partners’ dependence at the individual level is consistent with previous findings at the species level (Jordano 1987; Bascompte *et al*. 2006; Guimarães *et al*. 2006; Guerra & Pizo 2014). In Herrera (1984), most observed dependencies between frugivores and plant species were also weak or highly asymmetric. Interestingly, *P. lentiscus* showed quite symmetric dependencies –at the species level– with its main seed dispersers. Our analysis at the individual plant level revealed that, while these birds rely heavily on *P. lentiscus* fruits, they did not depend on particular plants but rather spread their dependencies, generating highly asymmetric interactions. If individual birds could have been identified too –rather than aggregated to species level– many of those plant strong dependencies on the main consumers might in turn transform into weak links, with just a few strong interactions (e.g., individual, territorial birds strongly depending on a specific patch of *P. lentiscus*). Hence, stepping down to the individual level seems important as it may enrich our perceptions of the embedded dependencies in mutualistic systems (Tonos et al. 2022) and address the proper scale to understand emerging properties at the species-level interaction networks (see Clark *et al*. 2011).

The available evidence suggests that symmetric dependencies could be rare in mutualistic systems. In fact, so far they have been reported only in very specific local communities, such as honeyeater-mistletoe facultative interactions (Reid 1990) or impoverished island systems (González-Castro *et al*. 2022). The disassortativity in the way species interact seems to promote asymmetry in partner’s dependence. Plants with a low degree (i.e., visited by one or few species) interacted with super-generalist avian species *C. melanocephala* and *E. rubecula*, which were also the most effective dispersers. At the same time, rare birds depended mostly on generalist plants. This favoured absence of symmetry in the dependence of rare species, agreeing with previous work arguing that reciprocal specialisations are rare (Joppa *et al*. 2009).

### Concluding remarks

Interactions between the individuals of a super-generalist plant with its fruit consumers have shown to be highly reciprocal in terms of the exchange of their mutualistic service, despite partners being highly asymmetric in their mutual dependence. These aspects appear quite general to low-intimacy mutualisms among free-living species (e.g., pollination, seed dispersal) which are largely dependent upon interaction frequency for the harvesting of food resources by animals. A key feature for the great success of super-generalists organisms appears to be related to abundance parameters that define their interaction frequency and, ultimately, their fitness. In contrast, highly specialised interactions likely depend on the ability to maintain reciprocity by means of a fine-tuned quality service between interacting species, where dependencies between partners would likely be more symmetric and intimate (Guimarães et al. 2007, Kiers et al. 2011). We might expect the emergence of these patterns when mutualisms among free-living species rely on encounter frequencies whose variance among species is so large as to obscure variation in the quality of outcome. Exceptions may include some mutualisms in specific environmental settings (e.g., oceanic islands) or characterised by high specificity of the interaction. Further studies on the reward reciprocity of generalised mutualistic interactions will build up more evidence to better understand the compromise between animals and plants in these mutualisms and the mechanisms behind the perpetuation of mutually-beneficial relationships.

## ACKNOWLEDGEMENTS

We are grateful to Juan Miguel Arroyo who provided essential support in the laboratory by working on the DNA-barcoding analysis. Camera data collection was possible thanks to the help of Lewis Barrett and especially Antonio López-Orta who assisted with long-hours of video screening. Video screening time was greatly facilitated by the use of DeepMeerkat on the EBD-CSIC data server facility, thanks to the assistance of Ben Weinstein (software developer) on parameter selection and Luis Torres on software installation. EQ is grateful to Rosario and Luis Carlos who kindly helped by measuring individual plants’ fruit attributes. Discussions with Jorge Isla, Blanca Arroyo and Luísa Genes were invaluable to help improve and elucidate the implications of this study. We thank the logistic and facilities support form ICTS-RBD Doñana and the Doñana National Park for onsite access authorizations during the fieldwork.

## Funding

The project leading to these results has received funding from “la Caixa” Foundation (ID 100010434), under agreement LCF/BQ/DE18/11670007. Authors were supported by project CGL2017-82847-P from the Agencia Estatal de Investigación, Spain and a LifeWatch ERIC-SUMHAL project (LIFEWATCH-2019-09-CSIC-13) with FEDER-EU funding. FRS was supported by the Talent Attraction programme from the VI Plan Propio de Investigación at Universidad de Sevilla (VI PPIT-US-2018-IV.2) and grant US-1381388 funded by FEDER 2014-2020 and Consejería de Economía, Conocimiento, Empresas y Universidad of Junta de Andalucía.

## Notes

### Competing Interest Statement

The authors have declared no competing interest.

