## Supplementary Material for "Reciprocity and interaction effectiveness in generalised mutualisms among free-living species"

A. Plant populations and frugivore species 1
B. Plant-animal interaction frequency 3
B.1. DNA-barcoding sampling 3
B.2. Camera-trap sampling 4
B.3. Interaction accumulation curves 7
C. Interaction outcome for birds (quality of resource provisioning effectiveness) 10
D. Interaction outcome for plants (quality of seed dispersal effectiveness) 11
D.1. Seeds escaping avian predation 11
D.2. Microhabitat seed deposition 11
D.3. Seeds escaping rodent predation 12
D.4. Seedling emergence and survival 14
E. Effectiveness calculations 16
E.1. Quantity component 17
E.2. Quality component of Resource Provisioning Effectiveness 29
E.3. Quality component of Seed Dispersal Effectiveness 30
E.4. Complete SDE landscape (including non-legitimate dispersers) 37
E.5. Variance partitioning of effectiveness components 38
F. Reciprocity and Asymmetry calculations 39
F.1. Analysis of reciprocity 39
F.2. Dependence and asymmetry calculations 40
G. Effects on consumption (quantity component) 42
G.1. Proportion of plants' crop consumed 42
G.2. Birds consumption of available energy 43
G.3. Predictors of fruit consumption intensity from individual plants 44
H. Null models for interaction asymmetry estimates 45
I. Software 47
J. References 50

### A. Plant populations and frugivore species

We sampled two study sites in Doñana National Park (Huelva, Spain): La Mancha del Rabicano in El Puntal site (EP; coords: 36.965180, -6.446582) and Laguna de las Madroñas (LM; coords: 37.030317, -6.471945). Both areas consist of Mediterranean sclerophyllous scrubland dominated by lentiscs (*Pistacia lentiscus)* coexisting with other fleshy-fruited species such as *Phillyrea angustifolia*, *Olea europaea* var. *sylvestris*, *Asparagus aphyllus* and *Myrtus communis.* The presence of pine trees (*Pinus pinea*) is scattered at EP, but more abundant at LM. The lower sclerophyllous scrubland is dominated by *Ulex parviflorus*, *Halimium halimifolium* and *Cistus salviifolius*. We used 2-4 ha plots within more extensive areas (over ca. 50 ha) of *P. lentiscus*-dominated shrubland, being surrounded by successional low shrubland dominated by *Halimium halimifolium* in drier places and *Erica arborea* in more humid locations (Allier *et al.* 1974; Rivas-Martínez *et al.* 1980).

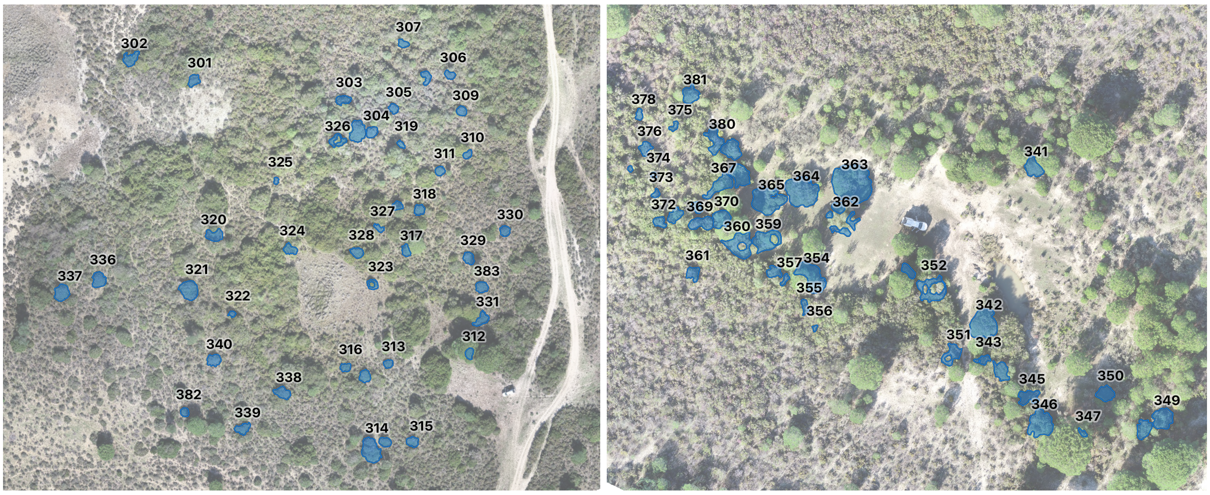

**Figure S.A.1**. Aerial image showing individual plants of *Pistacia lentiscus* marked at El Puntal (EP) and Laguna de las Madroñas (LM) populations; 40 plants per study site. The individual plants' canopies are outlined in blue and numbered.

*P. lentiscus* fruits are a staple food for frugivorous birds. Both the unripe (red) and ripe (black) fruits often have empty seeds as a result of either parthenocarpy, embryo abortion or pre-dispersal seed predation (Grundwag 1976; Jordano 1989). Frugivores strongly prefer the black, ripe fruits, and these typically have a higher proportion of filled, viable seeds (Jordano 1988, 1989) yet they also consume (in lower proportion) red fruits, which frequently have empty seeds. As a result frugivores mostly disperse filled, viable seeds but together with a variable fraction of empty seeds (González-Varo et al. 2019). The frequency of empty seeds varies greatly from year to year, as well as among *P. lentiscus* populations (Jordano 1988, 1989; Verdú

& García-Fayos 1998), resulting in variable amounts of empty seeds in the seed rain. In the focal study population, the mean percentage of empty seeds found in the plant canopy (estimated by floatability) was 67.5% at EP and 64.8% at LM (± 20.6% and 24.9% of SD respectively). At each site, we monitored 40 individual *P. lentiscus* plants for the complete 2019-20 fruiting season, totaling 80 focal individuals (Fig. S.A.1).

**Table S.A.1**. Frugivorous avian species considered in the study, average body mass, type of fruit consumption, and migratory status in the area. Types of frugivory acronyms: SD, seed disperser; SP, seed predator; PC, pulp consumer; PC/SD, pulp consumer with sporadic legitimate dispersal of seeds. Species are ordered by body mass (from Wilman *et al.* 2014).

| Species | Body mass (g) | Type of frugivory | Migration |
|---|---|---|---|
| *Columba palumbus* | 490.00 | SD/SP | Resident |
| *Turdus viscivorus* | 117.37 | SD | Winter migrant |
| *Turdus merula* | 102.73 | SD | Resident |
| *Cyanopica cooki* | 95.91 | SD | Resident |
| *Sturnus unicolor* | 83.66 | SD | Resident |
| *Turdus philomelos* | 67.74 | SD | Winter migrant |
| *Lanius meridionalis* | 60.43 | SD | Resident |
| *Coccothraustes coccothraustes* | 56.63 | SP | Winter migrant |
| *Chloris chloris* | 26.00 | SP | Resident |
| *Pyrrhula pyrrhula* | 24.26 | PC/SD | Winter migrant |
| *Fringilla coelebs* | 23.81 | PC/SD | Resident |
| *Curruca hortensis* | 21.90 | SD | Summer migrant |
| *Luscinia megarhynchos* | 19.60 | SD | Summer migrant |
| *Sylvia borin* | 18.20 | SD | Summer migrant |
| *Erithacus rubecula* | 17.70 | SD | Winter migrant |
| *Sylvia atricapilla* | 16.70 | SD | Winter migrant |
| *Parus major* | 16.25 | PC/SD | Resident |
| *Muscicapa striata* | 15.90 | SD | Summer migrant |
| *Curruca communis* | 15.10 | SD | Summer migrant |
| *Phoenicurus phoenicurus* | 14.59 | SD | Summer migrant |
| *Saxicola rubicola* | 14.09 | SD | Resident |
| *Ficedula hypoleuca* | 13.79 | SD | Summer migrant |
| *Cyanistes caeruleus* | 13.30 | PC/SD | Resident |
| *Curruca melanocephala* | 11.70 | SD | Resident |
| *Hippolais polyglotta* | 11.00 | SD | Summer migrant |
| *Curruca undata* | 10.80 | SD | Resident |
| *Curruca cantillans* | 9.60 | SD | Summer migrant |

### B. Plant-animal interaction frequency

We used two distinct sampling methods to monitor interaction frequency of frugivores and plants: DNA-barcoding of bird faecal and regurgitated samples and continuous-monitoring cameras.

*B.1. DNA-barcoding sampling*

Seed traps of 55 x 40 cm (0.22 $m^2$ trays) were located beneath the crown of individual plants, protected by a mesh of 1cm to prevent rodent predation. We placed one tray beneath every plant, except in four very large plants where we placed two trays. Seed traps were scanned fortnightly and all regurgitated and faecal samples in the tray were collected, regardless if they contained seeds or not. A total of 2691 samples were collected (1913 for EP and 778 for LM). On a few occasions, when the samples found in the trays were very abundant and presented identical aspect (e.g., multiple regurgitated seeds below a perch), a subset of samples were collected and the remaining count of seeds was assigned to the same species identified in the subset of samples obtained. Samples imputed this way represent 8% of the total samples obtained.

Animal-origin DNA was obtained from the surface of the samples (either scats or regurgitated seeds) was extracted and amplified using the primers COI-fsdF and COI-fsdR that target the COI region (cytochrome C oxidase subunit I; see González-Varo *et al.* 2014). Amplified DNA was then sequenced and identified using the Barcode Of Life Data (BOLD) Systems database (https://www.boldsystems.org/) or the Nucleotide Basic Local Alignment Search Tool (BLAST) from the NCBI (https://blast.ncbi.nlm.nih.gov/Blast.cgi). DNA-barcoding analysis was carried out following the protocol described in González-Varo *et al.* (2014) with some modifications. In order to reduce time and costs, silica suspension addition step was removed, where instead DNA supernatant and binding buffer were added directly to the column with the microfiber filter. The column was then set for the second incubation period. This modification was based on the finding that DNA similarly attaches to the glass microfiber filter (Shi *et al.* 2018). Replacing silica suspension by glass microfiber filter we obtained similar identification yields and successful amplification rates. Columns brand (MoBiTec, Germany) was also replaced by another brand (Sarstedt, Germany).

The number of samples collected under some plants was considerably high (over 200 samples in some cases). To ensure that the assemblage of avian visitors to individual plants was well characterised, we proceeded with DNA-barcoding laboratory analysis until sampling completeness was reasonably robust. A minimum of 40 samples per plant were analysed. This minimum however was subjected to sample availability, as some individuals had few samples or these were highly degraded samples not suitable for analysis. For plants with more than 40 samples, we gradually increased the number of DNA-barcoded samples until the sampling completeness curves were saturated (see Suppl. Mat S.B.3). In cases where individuals had fewer than 40 samples, we processed all the available samples for each individual that were suitable for analysis (i.e. removing samples with highly degraded DNA). In total, we analysed 90% and 96% of the samples collected for EP and LM respectively. Identification success rate of the analysed samples was 94% (n = 2285). We established a quality criterion for DNA-barcoded samples, where we only considered samples over 150 bp length and over 90% of identity similarity. Most samples, however, scored over 99% similarity (mean length = 288 bp, mean similarity = 99.31%). For the minority subset of samples whose similarity was between 90%-99% (n = 228), the second species identified had to be further than 2% similarity distance, or absent in the geographical range area, as an additional quality requisite.

*B.2. Camera-trap sampling*

In addition to DNA barcoding, we also used video monitoring to record animal visitation and feeding events in focal plants at EP site. Continuous-monitoring cameras (GoPro Hero® 7 White) were set facing individual plants, so that almost all of the plant could be seen from one side (Fig. S.B.2.1). We recorded plants nine times spaced along the season, however some differences may exist between total recording times due to camera issues (see Table S.B.2.1). Just in a few occasions, cameras turned off earlier due to battery issues or SD card was illegible. Cameras started recording  between 8:00-10:00am for a period of approximately 2.2 hours. All individual plants were monitored every fortnight for a total of 9 times along the fruiting season, accumulating more than 19 hours of observation per individual plant on average (range = 18-20). Overall, cameras recorded 3790 visits by avian frugivores.

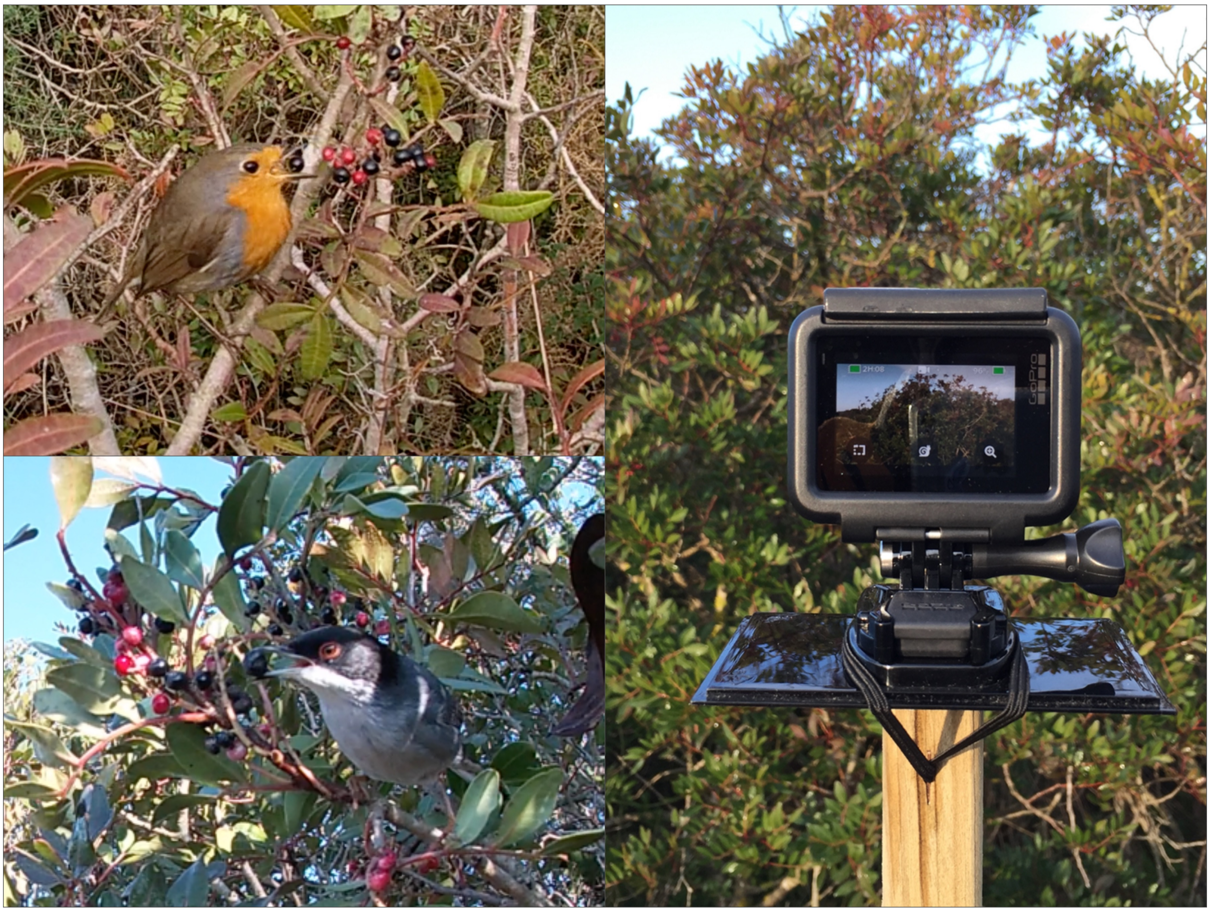

**Figure S.B.2.1**. Photos captured from GoPro video recordings of *Erithacus rubecula* (top left) and *Curruca melanocephala* (bottom left) feeding on *Pistacia lentiscus* fruits. The image on the right shows the installation of the GoPro camera to record the interactions of visiting birds on individual plants.

We analysed the >700 hours of video recordings with the help of the motion detection program DeepMeerkat (Weinstein 2018). Motion detection helped to locate the specific moment of a visitation event, narrowing considerably the video screening time for analysis. DeepMeerkat was most helpful when the wind was mild, otherwise there were too many false positives caused by moving branches; in these cases the videos were fully watched to detect visitation events. We performed several trials to determine the best parameter threshold at which the DeepMeerkat algorithm was most sensitive (*i.e.* detected most true positives), and settled on a tensorflow threshold (i.e., confidence level to ignore movement detected; Weinstein 2018) of $10^{-11}$ and a minimum size of contour of $10^{-10}$. We also carried out a parallel analysis of 22 videos to test the success rate of DeepMeerkat motion detection in comparison with detection by naked eye. Of a total of 46 interactions recorded in the test videos, four interactions were exclusively detected by DeepMeerkat and three by the naked eye, indicating very good performance of DeepMeerkat even though there is some trade-off between both approaches. The species exclusively detected with the program were mainly perching species (*P. phoenicurus* and *F.*

*hypoleuca*) that can pass undetected to the naked eye by their stillness, whereas their fast arrival can be detected by the program. On the other hand, the three naked eye exclusive detections corresponded to *Curruca melanocephala* that tends to scurry around the plant, being easier to detect by the naked eye in a fast-paced video, but may become more cryptic for the program if the animal is moving behind vegetation.

For every visitation event we recorded the identity of the visiting species when possible, arrival and departure time, visit length, behaviour and number of fruits consumed, if any. Species identification was possible for 91% of the visits (n = 323 visits by unknown species). We extracted information on the feeding frequency of animal species (i.e. fraction of visits where there was actual fruit consumption) and the number of fruits consumed per visit. We could detect feeding on fruits and/or seeds on 927 out of 3790 visits (24%), and recorded the number of consumed fruits or seeds whenever possible. A total of 37 animal species were identified to be interacting with the individual plants, of which 26 species were frugivorous birds (species known to feed on *P. lentiscus* fruits, even sporadically).

**Table S.B.2.1.** Time in minutes spent recording individual plants along nine different periods.

| ID plant | Sept. | Sept. | Oct. | Oct. | Nov. | Nov. | Dec. | Dec. | Jan. | Total time (min) |
|---|---|---|---|---|---|---|---|---|---|---|
| **301** | 94 | 136 | 136 | 136 | 136 | 76 | 135 | 135 | 133 | 1117 |
| **302** | 88 | 135 | 136 | 136 | 136 | 136 | 136 | 135 | 135 | 1173 |
| **303** | 124 | 113 | 136 | 136 | 136 | 136 | 135 | 135 | 135 | 1186 |
| **304** | 96 | 120 | 136 | 136 | 136 | 136 | 135 | 126 | 135 | 1156 |
| **305** | 114 | 128 | 130 | 136 | 130 | 136 | 98 | 125 | 135 | 1132 |
| **306** | 127 | 135 | 136 | 136 | 127 | 136 | 135 | 121 | 135 | 1188 |
| **307** | 135 | 136 | 136 | 135 | 129 | 136 | 135 | 124 | 136 | 1202 |
| **308** | 35 | 135 | 136 | 135 | 128 | 136 | 135 | 121 | 135 | 1096 |
| **309** | 134 | 122 | 119 | 136 | 136 | 136 | 135 | 135 | 73 | 1126 |
| **310** | 122 | 124 | 121 | 136 | 136 | 136 | 136 | 135 | 134 | 1180 |
| **311** | 120 | 126 | 79 | 136 | 136 | 136 | 135 | 135 | 74 | 1077 |
| **312** | 131 | 130 | 130 | 136 | 109 | 136 | 136 | 135 | 135 | 1178 |
| **313** | 131 | 104 | 114 | 136 | 136 | 136 | 130 | 135 | 135 | 1157 |
| **314** | 131 | 136 | 136 | 134 | 136 | 136 | 135 | 135 | 133 | 1212 |
| **315** | 110 | 110 | 136 | 136 | 136 | 120 | 135 | 135 | 135 | 1153 |
| **316** | 115 | 135 | 136 | 136 | 136 | 136 | 135 | 135 | 135 | 1199 |
| **317** | 134 | 103 | 136 | 136 | 136 | 136 | 135 | 135 | 135 | 1186 |
| **318** | 135 | 129 | 136 | 135 | 135 | 136 | 135 | 135 | 135 | 1211 |
| **319** | 52 | 124 | 131 | 135 | 131 | 136 | 135 | 125 | 135 | 1104 |
| **320** | 112 | 135 | 133 | 136 | 135 | 136 | 135 | 135 | 135 | 1192 |
| **321** | 118 | 18 | 136 | 136 | 136 | 136 | 135 | 135 | 131 | 1081 |
| **322** | 0 | 135 | 136 | 136 | 136 | 136 | 135 | 135 | 127 | 1076 |
| **323** | 135 | 134 | 136 | 136 | 136 | 136 | 135 | 136 | 135 | 1219 |

| | | | | | | | | | | |
|---|---|---|---|---|---|---|---|---|---|---|
| **324** | 135 | 118 | 136 | 136 | 136 | 136 | 135 | 135 | 135 | 1202 |
| **325** | 131 | 125 | 136 | 136 | 136 | 136 | 136 | 135 | 135 | 1206 |
| **326** | 124 | 105 | 136 | 135 | 133 | 136 | 135 | 127 | 135 | 1166 |
| **327** | 130 | 131 | 136 | 136 | 136 | 136 | 135 | 135 | 135 | 1210 |
| **329** | 100 | 88 | 123 | 136 | 136 | 136 | 135 | 135 | 47 | 1036 |
| **330** | 133 | 126 | 122 | 136 | 116 | 136 | 135 | 136 | 112 | 1152 |
| **331** | 135 | 128 | 128 | 136 | 136 | 136 | 135 | 135 | 49 | 1118 |
| **332** | 81 | 130 | 136 | 135 | 134 | 136 | 135 | 134 | 136 | 1157 |
| **334** | 50 | 119 | 136 | 136 | 100 | 136 | 135 | 135 | 135 | 1082 |
| **335** | 131 | 136 | 57 | 136 | 136 | 136 | 123 | 135 | 135 | 1125 |
| **336** | 86 | 119 | 136 | 136 | 136 | 136 | 135 | 135 | 135 | 1154 |
| **337** | 127 | 136 | 136 | 136 | 136 | 136 | 135 | 135 | 135 | 1212 |
| **338** | 118 | 136 | 136 | 136 | 136 | 135 | 136 | 90 | 132 | 1155 |
| **339** | 206 | 136 | 88 | 136 | 136 | 136 | 135 | 135 | 47 | 1155 |
| **340** | 0 | 136 | 136 | 136 | 136 | 135 | 135 | 135 | 134 | 1083 |
| **382** | 123 | 134 | 136 | 136 | 136 | 136 | 49 | 135 | 132 | 1117 |
| **383** | 87 | 127 | 125 | 136 | 136 | 136 | 135 | 135 | 75 | 1092 |

*B.3. Interaction accumulation curves*

We used interaction accumulation curves (IAC, analogous to species accumulation curves) to determine both DNA-barcoding and video recording sampling completeness (Colwell & Coddington 1994; Jordano 2016). The number of samples collected in seed traps under individual lentiscs varied from 2 up to 203 for the whole fruiting season. Most plants (72 out of 80) had up to 90% of their samples analysed (see Fig. S.B.3.1, Table S.B.3.2). Overall sampling completeness was 93% for both methods (sensu Chacoff *et al.* 2012); 95% for cameras and 96% for DNA-barcoding (Table S.B.3.1). The total number of frugivorous species recorded was 27; of which 26 were recorded with cameras and 22 with DNA-barcoding.

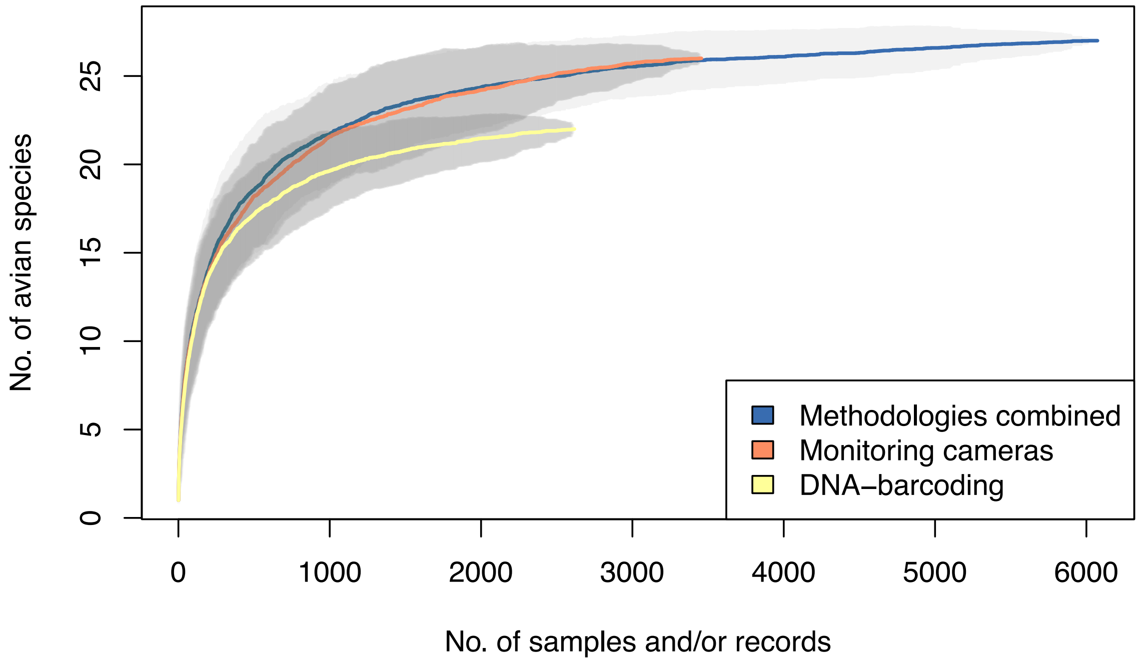

**Figure S.B.3.1**. Interaction Accumulation Curves for animal interaction records with *Pistacia lentiscus* using two different methodologies, and the result of considering both together by combining the two datasets (see Quintero et al. 2021).

**Table S.B.3.1.** Sampling completeness for interactions with *Pistacia lentiscus* using datasets resulting from different methodologies. Number of samples, number of species recorded, Chao estimator and its Standard Error, as well as completeness (sensu Chacoff *et al.* 2012) are provided. For the DNA barcoding samplings, results are given for EP and LM sites separately. Note that in LM site the Chao estimator is rather uncertain (SE > 10), hence the estimated completeness is rather uncertain too; in all cases, however, the empirical values fall within the ±1SE range of the estimator.

| Dataset | N samples | Species | Chao | Chao SE | Completeness |
|---|---|---|---|---|---|
| Both methodologies | 6073 | 27 | 29.0 | 3.7 | 0.93 |
| Monitoring cameras | 3456 | 26 | 27.5 | 2.3 | 0.95 |
| DNA-Barcoding | 2617 | 22 | 23.0 | 2.3 | 0.96 |
| DNA-Barcoding EP | 1851 | 21 | 22.0 | 1.9 | 0.95 |
| DNA-Barcoding LM | 766 | 16 | 26.0 | 10.2 | 0.62 |

**Table S.B.3.2.** Number of species recorded per individual plant, total number of correctly identified frugivorous interactions records and its breakdown by the methodologies used. The total number of records refers to all samples/videos considered in the study (considering only successfully identified DNA-barcoding samples and only from avian frugivores). The table also shows the number of samples collected and analysed for DNA-barcoding methodology, as well as the total number of visits recorded and identified for the camera-traps. 'P' indicates the proportion of samples/videos analysed/identified with each methodology relative to the total obtained for each plant. Results are given separately for El Puntal (EP) and Las Madroñas (LM) sites, with only the former being monitored with the two methods.

El Puntal

| | | | DNA-barcoding | | | Cameras | | |
|---|---|---|---|---|---|---|---|---|
| **Plant ID** | **spp** | **Total records** | Samples collected | Samples analysed | P | Visits recorded | Visits identified | P |
| 301 | 11 | 214 | 40 | 38 | 0.95 | 189 | 181 | 0.96 |
| 302 | 15 | 196 | 27 | 27 | 1.00 | 194 | 172 | 0.89 |
| 303 | 9 | 68 | 21 | 21 | 1.00 | 50 | 48 | 0.96 |
| 304 | 19 | 465 | 203 | 118 | 0.58 | 392 | 350 | 0.89 |
| 305 | 5 | 40 | 20 | 18 | 0.90 | 25 | 22 | 0.88 |
| 306 | 11 | 142 | 54 | 52 | 0.96 | 105 | 98 | 0.93 |
| 307 | 7 | 61 | 23 | 22 | 0.96 | 45 | 41 | 0.91 |
| 308 | 7 | 134 | 71 | 71 | 1.00 | 68 | 63 | 0.93 |
| 309 | 9 | 80 | 35 | 35 | 1.00 | 45 | 45 | 1.00 |
| 310 | 7 | 87 | 30 | 29 | 0.97 | 62 | 58 | 0.94 |
| 311 | 8 | 56 | 23 | 22 | 0.96 | 36 | 36 | 1.00 |
| 312 | 7 | 88 | 64 | 60 | 0.94 | 32 | 29 | 0.91 |
| 313 | 9 | 122 | 44 | 43 | 0.98 | 83 | 80 | 0.96 |
| 314 | 18 | 307 | 136 | 102 | 0.75 | 240 | 205 | 0.85 |
| 315 | 10 | 147 | 51 | 50 | 0.98 | 119 | 101 | 0.85 |
| 316 | 6 | 46 | 15 | 15 | 1.00 | 33 | 31 | 0.94 |
| 317 | 5 | 60 | 28 | 27 | 0.96 | 38 | 34 | 0.89 |
| 318 | 9 | 213 | 106 | 92 | 0.87 | 139 | 125 | 0.90 |
| 319 | 11 | 122 | 43 | 43 | 1.00 | 84 | 83 | 0.99 |
| 320 | 9 | 208 | 134 | 113 | 0.84 | 109 | 102 | 0.94 |
| 321 | 11 | 167 | 65 | 63 | 0.97 | 120 | 108 | 0.90 |
| 322 | 12 | 110 | 67 | 64 | 0.96 | 53 | 49 | 0.92 |
| 323 | 11 | 160 | 61 | 50 | 0.82 | 116 | 113 | 0.97 |
| 324 | 8 | 71 | 22 | 22 | 1.00 | 56 | 49 | 0.88 |
| 325 | 8 | 91 | 41 | 41 | 1.00 | 54 | 52 | 0.96 |
| 326 | 11 | 120 | 57 | 57 | 1.00 | 74 | 66 | 0.89 |
| 327 | 10 | 126 | 79 | 75 | 0.95 | 66 | 55 | 0.83 |
| 329 | 11 | 124 | 45 | 43 | 0.96 | 85 | 82 | 0.96 |
| 330 | 9 | 94 | 33 | 32 | 0.97 | 66 | 63 | 0.95 |
| 331 | 8 | 91 | 35 | 35 | 1.00 | 63 | 59 | 0.94 |
| 332 | 11 | 146 | 52 | 50 | 0.96 | 109 | 97 | 0.89 |
| 334 | 10 | 139 | 53 | 52 | 0.98 | 99 | 91 | 0.92 |
| 335 | 8 | 98 | 41 | 41 | 1.00 | 60 | 58 | 0.97 |
| 336 | 11 | 147 | 51 | 51 | 1.00 | 108 | 99 | 0.92 |
| 337 | 17 | 244 | 65 | 64 | 0.98 | 205 | 183 | 0.89 |
| 338 | 12 | 206 | 94 | 84 | 0.89 | 146 | 125 | 0.86 |
| 339 | 7 | 42 | 16 | 16 | 1.00 | 28 | 26 | 0.93 |
| 340 | 10 | 120 | 28 | 28 | 1.00 | 104 | 94 | 0.90 |
| 382 | 8 | 37 | 19 | 19 | 1.00 | 20 | 18 | 0.90 |
| 383 | 7 | 118 | 55 | 55 | 1.00 | 70 | 65 | 0.93 |

Madroñas

| | | | DNA-barcoding | | |
|---|---|---|---|---|---|
| **Plant ID** | **spp** | **Total records** | Samples collected | Samples analysed | P |
| 341 | 3 | 22 | 22 | 22 | 1.00 |
| 342 | 5 | 17 | 19 | 19 | 1.00 |
| 343 | 3 | 24 | 24 | 24 | 1.00 |
| 344 | 5 | 18 | 19 | 19 | 1.00 |
| 345 | 3 | 5 | 8 | 8 | 1.00 |
| 346 | 4 | 45 | 46 | 46 | 1.00 |
| 347 | 5 | 18 | 18 | 18 | 1.00 |
| 348 | 4 | 39 | 39 | 39 | 1.00 |
| 349 | 2 | 10 | 10 | 10 | 1.00 |
| 350 | 5 | 46 | 53 | 48 | 0.91 |
| 351 | 4 | 7 | 8 | 8 | 1.00 |
| 352 | 4 | 47 | 56 | 48 | 0.86 |
| 353 | 5 | 21 | 21 | 21 | 1.00 |
| 354 | 3 | 34 | 34 | 34 | 1.00 |
| 355 | 3 | 12 | 13 | 12 | 0.92 |
| 356 | 1 | 9 | 9 | 9 | 1.00 |
| 357 | 3 | 11 | 11 | 11 | 1.00 |
| 358 | 3 | 16 | 17 | 17 | 1.00 |
| 359 | 3 | 12 | 12 | 12 | 1.00 |
| 360 | 4 | 12 | 13 | 13 | 1.00 |
| 361 | 2 | 11 | 11 | 11 | 1.00 |
| 362 | 2 | 6 | 14 | 14 | 1.00 |
| 363 | 3 | 29 | 32 | 32 | 1.00 |
| 364 | 6 | 38 | 56 | 40 | 0.71 |
| 365 | 4 | 25 | 32 | 31 | 0.97 |
| 366 | 3 | 26 | 26 | 26 | 1.00 |
| 367 | 3 | 19 | 23 | 21 | 0.91 |
| 368 | 3 | 10 | 10 | 10 | 1.00 |
| 369 | 3 | 11 | 13 | 13 | 1.00 |
| 370 | 4 | 9 | 10 | 10 | 1.00 |
| 371 | 1 | 12 | 12 | 12 | 1.00 |
| 372 | 2 | 8 | 9 | 9 | 1.00 |
| 373 | 3 | 11 | 12 | 12 | 1.00 |
| 374 | 1 | 1 | 2 | 2 | 1.00 |
| 375 | 3 | 11 | 12 | 12 | 1.00 |
| 376 | 3 | 11 | 12 | 12 | 1.00 |
| 378 | 2 | 14 | 14 | 14 | 1.00 |
| 379 | 5 | 47 | 51 | 49 | 0.96 |
| 380 | 3 | 29 | 30 | 30 | 1.00 |
| 381 | 6 | 13 | 14 | 14 | 1.00 |

### C. Interaction outcome for birds (quality of resource provisioning effectiveness)

To estimate differences in fruit quality provided by individual plants, we randomly collected ripe fruits (mean = 31 fruits, range = 17-63) from each individual plant at both populations and measured the whole fruit and the seed fresh mass. Pulp mass was calculated as the difference in weight between the whole fruit and the seed, i.e., before and after being manually depulped (Fig. S.C.1). This pulp and seed mass was later converted into energy obtained (see Suppl. Mat. E.2), depending on the bird feeding behaviour (frugivorous or granivorous; see Table S.A.1. for bird species categorization into frugivory types).

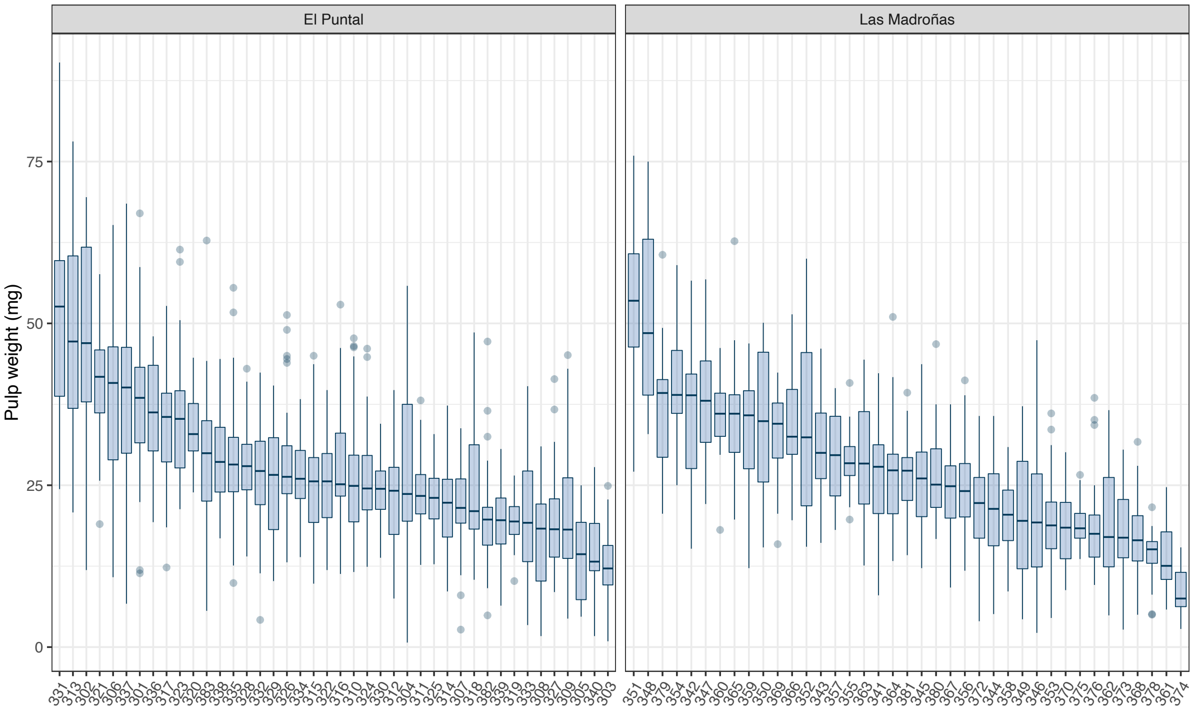

**Figure S.C.1.** Pulp fresh mass per fruit (in mg) for individual plants in the two study populations. Each box represents the 1st-3rd interquartile range, the solid middle line represents the median, and whiskers extend to the largest or smallest value no further than 1.5 times the interquartile range; dots indicate more extreme values. Numbers at the bottom are the individual plant identification codes.

### D. Interaction outcome for plants (quality of seed dispersal effectiveness)

In order to estimate the probabilities of seedling recruitment resulting from fruit/seed consumption by each avian consumer we considered four steps in chronological order: (1) probability of seeds to escape granivorous birds predation during handling, (2) microhabitat use patterns by each bird species, (3) probability of seeds escaping rodent post-dispersal predation in each microhabitat, and (4) probability of seedling emergence and early survival (past through their first summer) in each specific microhabitat.

*D.1. Seeds escaping avian predation*

Some intact clean seeds found in the seed traps were attributed to *Chloris chloris*, *Fringilla coelebs* and *Pyrrhula pyrrhula* through DNA-barcoding (n = 36). This indicates that sporadic dispersal events by these granivores are possible if intact seeds are dropped during handling. To take this into account, we calculated the probability of seeds escaping predation by avian granivores using the total number of preyed-upon seeds (open seed endocarp halves) and the number of intact seeds attributed to granivores found in each seed tray.

*D.2. Microhabitat seed deposition*

We classified the vegetation of both sites into five microhabitats for measuring seed dissemination and establishment success: (1) under *Pistacia lentiscus* conspecifics (PL), (2) under other fleshy fruited species (FR), (3) under non-fleshy fruited species (NF), (4) under pine trees (*Pinus pinea*; PP), and (5) open ground areas (OA). We expected different bird species to use these microhabitats with varying intensity, hence generating contrasting seed rain abundance and composition. Expected microhabitat variation in seed predator abundance and microclimatic conditions would also affect the fate of dispersed seeds (García *et al.* 2005; Gómez-Aparicio 2008).

In order to estimate the probability of dispersal of *Pistacia lentiscus* seeds towards each microhabitat, we collected dispersed seeds in the five microhabitats distributed along El Puntal (EP) area. For the *Pistacia lentiscus* (PL) microhabitat, we included all dispersed seeds collected in the seed trays beneath the 40 individual *P. lentiscus* plants monitored at EP site. For the other three microhabitats beneath vegetation cover (FR, NF and PP) we placed two seed-sampling trays (33 x 25.5 cm; 0.084 $m^2$) in 15 replicated locations per microhabitat. Lastly, for the open ground area (OA), we sampled 17 transects, 100 to 400 metres long and 1 m wide, at different times distributed along the fruiting season, and collected every faeces containing *P. lentiscus* seeds.

A total of 1664 seeds of *Pistacia lentiscus* were collected in the five microhabitats, of which 96% were analysed (n = 1594 seeds). The identity of the bird dispersing the seeds was determined through DNA-barcoding analysis, using the same protocol described above (see also Suppl. Mat. B.1). DNA-barcoding identification success was 95%. The number of *P. lentiscus* seeds dispersed by each bird species to each microhabitat were then used to estimate their differential contributions to seed rain across microhabitats (see below).

*D.3. Seeds escaping rodent predation*

To estimate post-dispersal predation rates we placed 6 experimental predation station replicates per microhabitat. Each experimental unit consisted of a petri dish open to rodents and a control plate protected with wire mesh of 1cm light to prevent rodent predation, each containing 10 seeds (Fig. S.D.3.1). These controls allowed us to discern when the disappearance of a seed was not caused by rodents but by other animals, most likely ants. All seeds were ensured to be viable through flotation-sink experiments (Albaladejo *et al.* 2009) to avoid empty seed detection by the animals (Jordano 1989). The experimental units were checked every one or few days at the beginning of the experiment and then checks were gradually spaced over time (Fig. S.D.3.2). Experimental units were installed in January 2019 and removed in July 2019, for a total of 131 days.

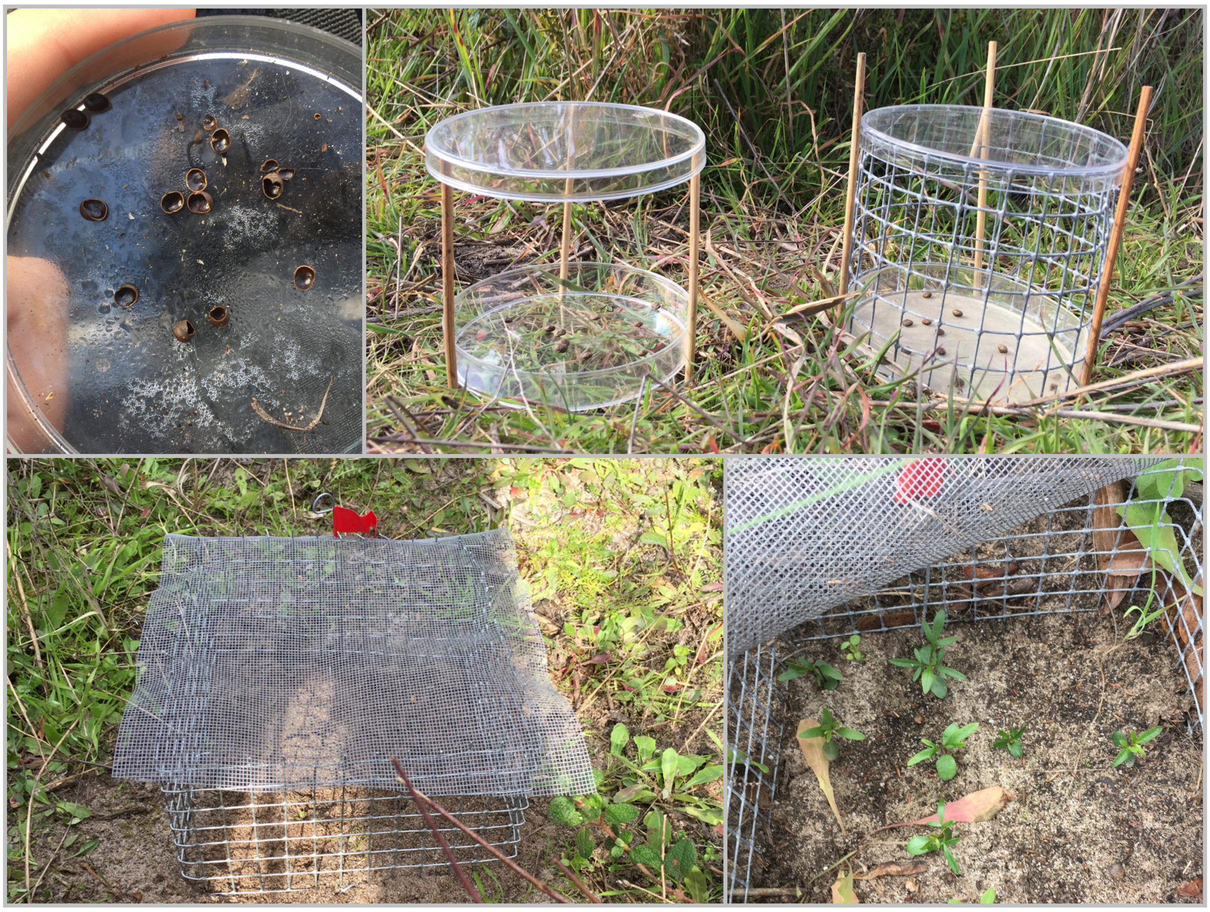

**Figure S.D.3.1.** Photos of (top left) broken seeds without embryo after being preyed upon by rodents (*Mus spretus* and/or *Apodemus sylvaticus*); (top right) control (open) and experimental (protected with mesh wire) seed predation stations used in the field; (bottom left) experimental sowing station used in the field, and two-month old emerged seedlings (bottom right).

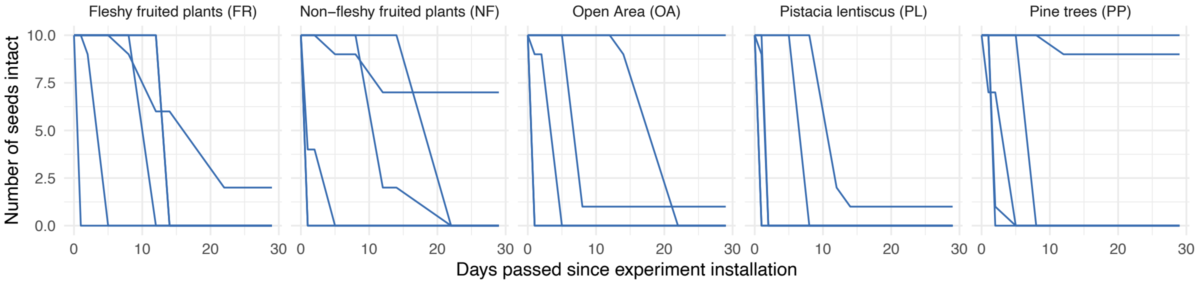

**Figure S.D.3.2.** Number of seeds surviving rodent predation in the five microhabitats along time. Each line corresponds to an experimental station, each starting with 10 intact viable seeds in the beginning. The dashed vertical line represents the 30-day cutoff, which we considered as the critical period for predation as seedlings start to emerge around the fourth week since sowing.

*D.4. Seedling emergence and survival*

We carried out experimental seed sowing to assess seedling emergence and early survival rates per microhabitat. These experiments were repeated for two consecutive years during the fruiting season of 2018-19 and 2019-20. We installed 6 to 7 replicates of germination stations in each microhabitat. Germination stations consisted of 16 sown seeds spaced 1.5 cm between each other in a four by four grid, and protected by a 1cm-light wire mesh on the sides and a fibreglass mesh on top to prevent herbivory, debris and trampling (Fig. S.D.3.1). All sowed seeds were checked to be viable through flotation-sink experiments and came from 8 and 6 different mothers for the first and second year, respectively. We ensured the mother origin of the seeds was equally distributed among all stations and microhabitats. Seeds were submerged in cold water for 24 h previous to sowing, as seedling emergence is conditioned to abundant rain events (García-Fayos & Verdú 1998; Del Campo *et al.* 2014). Germination experiments started in January of 2019 and in October of 2019. Seedling emergence and survival were monitored approximately every fortnight for the first four months after sowing and monthly thereafter until no seedlings remained alive (Fig. S.D.4.1; S.D.4.2).

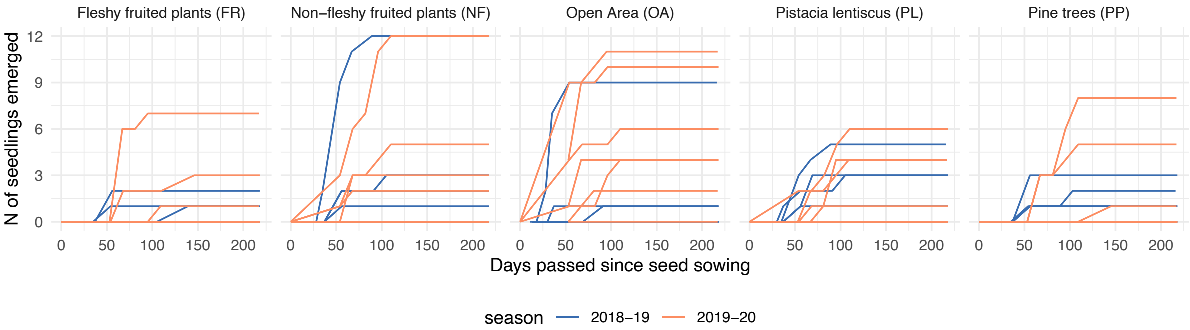

**Figure S.D.4.1.** Seedling emergence dynamics in the experimental sowing units set up across the five microhabitats in two different seasons (2018-19 and 2019-20). Each line represents an experimental unit, consisting of 16 seeds. The number of experimental units per microhabitat was 6 in 2018-19 and 7 in 2019-20.

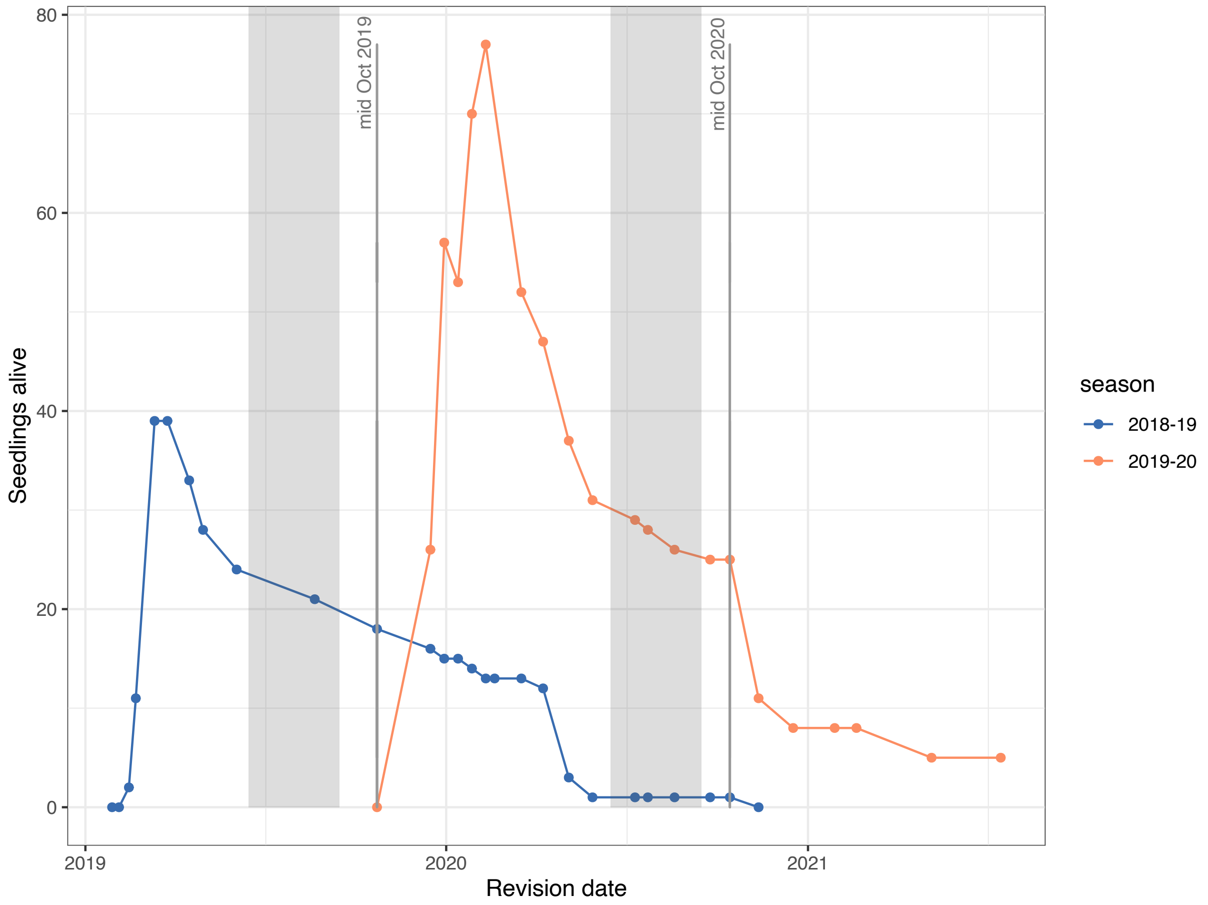

**Figure S.D.4.2.** Number of seedlings recorded alive in the experimental sowing stations during two consecutive fruiting seasons, from January 2019 to July 2021. The number of seedlings recorded in a given date includes newly emerged seedlings as well as those surviving from previous dates. The shaded area in grey corresponds to the hottest months (from 15th June to 15th of September). For each season we quantified seedling survival just after their first summer (mid October, grey vertical lines).

### E. Effectiveness calculations

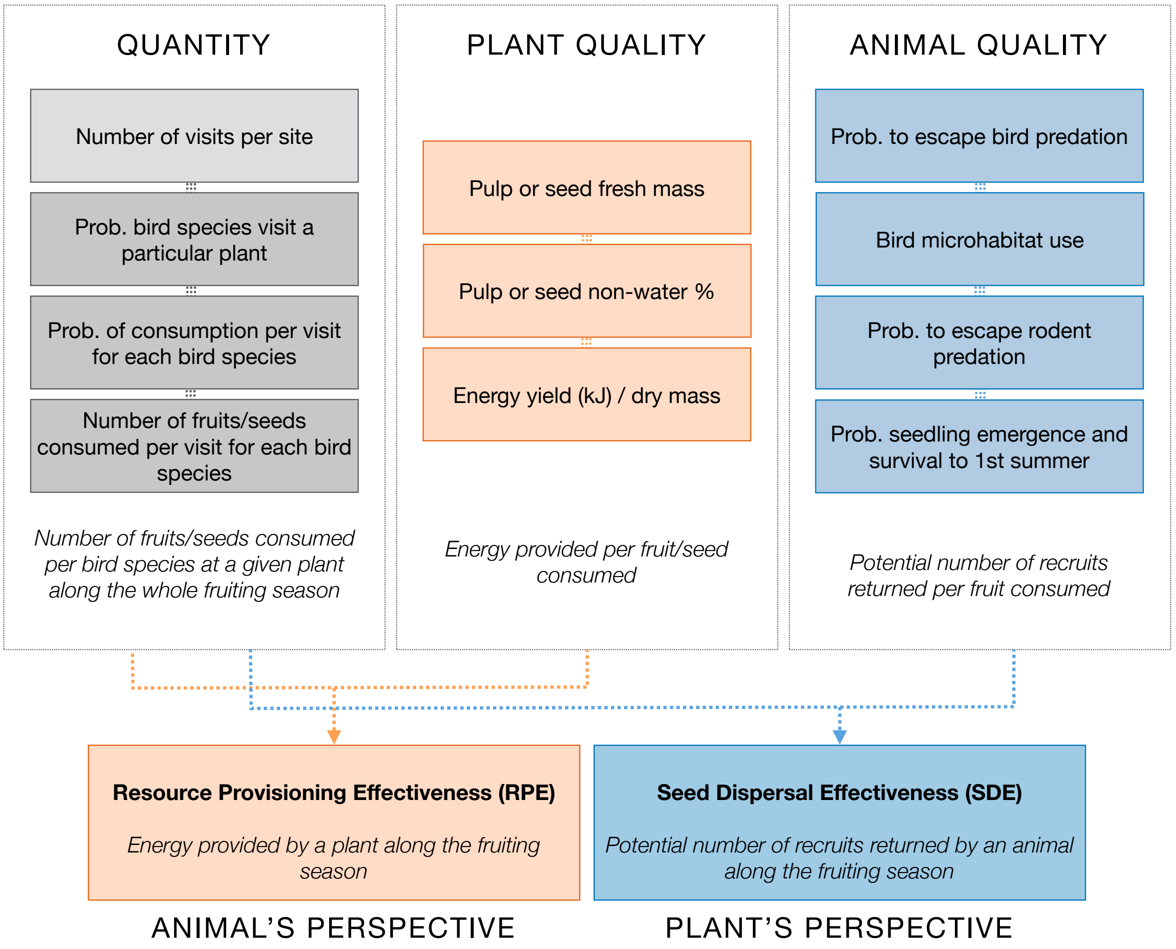

**Figure S.E.1.** Diagram of all the elements involved in SDE and RPE calculations. The estimates for each component (i.e. Quantity and Quality) are all chronological and multiplicative sequential steps (Schupp *et al.* 2017).

Calculating seed dispersal or resource provisioning effectiveness (SDE and RPE, respectively) requires large amounts of data (on bird visitation rates and fruit consumption patterns, seed rain density and post-dispersal survival, fruit weights, etc.; Fig. S.E.1) which are rarely available for all the plants and bird species involved. Most effectiveness studies try to fill data gaps *ad hoc*, e.g. assigning fruit consumption patterns from similar or related species, without considering uncertainties. Here we attempt a pure model-based approach to estimate all the components required to estimate SDE and RPE. In particular, we exploit hierarchical Bayesian models to share information ("borrow strength") across bird species and plant individuals, being able to obtain probabilistic estimates even for unobserved quantities. Furthermore, by using the full posterior distributions from all the estimated quantities we can propagate uncertainties and provide more realistic estimates of the uncertainty involved in such convoluted effectiveness analyses.

We explain how we estimated each component of RPE and SDE below. Once we had estimated each component, we multiplied the quantity and quality components to calculate the total effectiveness. The quantity component (i.e., total number of fruits consumed by a specific bird on a given plant) was common for both the animal and plant's perspective. Quality for the animal was the energy acquired per fruit/seed consumed. Resource Provisioning Effectiveness (RPE) therefore represents the total energy acquired by a specific bird along the fruiting season from a given plant. Quality for the plant was the probability that a consumed and viable fruit becomes a seedling surviving its first summer. Seed Dispersal Effectiveness therefore indicates the potential number of seedlings recruited for a specific plant by a given bird species.

*E.1. Quantity component*

We estimate the number of fruits consumed by each bird species at each individual plant combining the following quantities:

- Total number of bird visits received by plants at each site (estimated from bird droppings in seed traps beneath mother plants)
- Probability that different bird species visit a particular plant (estimated from both DNA barcoding and video cameras)
- Probability of fruit/seed consumption per visit for each bird species (estimated from video recordings)
- Number of fruits/seeds consumed per visit of each bird species (estimated from video recordings)

**Total number of visits and probability of visit to each individual plant by each bird species**

To estimate the probability of visit to each plant from each bird species we used data from DNA barcoding of droppings collected beneath mother plants (both sites), as well as data obtained through the analysis of video recordings (Puntal site only). Estimates from barcoding and video analysis for El Puntal site were then merged (see below).

*Estimating probability of visit from barcoding data*

For each of both sites, we used all bird droppings collected at seed trays beneath lentisc plants to estimate the total number of bird visits to each individual plant *i* along the fruiting season, using a hierarchical Bayesian Poisson regression:

$$Nvisit_{DNAi} \sim Poisson(\lambda_{DNAi})$$

where the log number of visits received by each individual *i* ($\lambda_{DNAi}$) was modelled as

$$log(\lambda_{DNAi}) = \mu_{visitDNA} + \alpha_{DNAi} + \text{offset}(log(\text{trap.area}_i))$$

$$\alpha_{DNAi} \sim N(0, \sigma^2_{DNAvisit})$$

In this equation, $\mu_{visitDNA}$ is the average number of bird visits across all individual plants over the season, and $\alpha_{DNAi}$ represents individual variation around that population average (i.e. a random intercept), drawn from a Normal distribution with standard deviation $\sigma_{DNAvisit}$. We included an offset term to account for the fact that sampling effort was not constant among individuals (4 plants at El Puntal site had 2 seed trays placed beneath, while all other individuals had 1 seed tray). Hence, all parameter estimates refer to visits/$m^2$ of canopy area. Note this model assumes that each bird dropping corresponds to a single visit.

We used weakly informative Normal priors for all parameters, and performed prior predictive checks in all models to ensure that our priors produced reasonable estimates. $\mu_{visitDNA}$ had a Normal(4, 1) prior in log scale, corresponding to c. 50 bird visits per square metre of canopy area over the whole season. $\sigma_{DNAvisit}$ had a half-Normal prior with standard deviation = 1, i.e. Normal(0, 1) truncated at 0.

Once we had estimated the number of visits to each individual plant (taking into account their total canopy area as measured from the drone image; Fig. S.A.1), we could calculate the total number of visits per site (aggregating all individuals) and the relative probability of visit of each individual plant ($Pvisit_{DNAi}$) by dividing their visits by the total number of visits at the site.

Then, we estimated the probability that a given visit is from a given bird species ($Pbird_{DNAij}$). In other words, the proportion of visits from each bird species (as identified from DNA barcoding) at each plant. For that, we modelled the number of visits from each bird species *j* to each individual plant *i* following a Binomial distribution and a logit link (*log(P/(1-P))*):

$$Nvisit_{DNAij} \sim Binomial(Nvisit_{DNAi}, Pbird_{DNAij})$$

$$logit(Pbird_{DNAij}) = \mu_{birdDNA} + \alpha_{DNAi} + \alpha_{DNAj} + \alpha_{DNAij}$$

$$\alpha_{DNAi} \sim N(0, \sigma^2_{DNAplant})$$

$$\alpha_{DNAj} \sim N(0, \sigma^2_{DNAbird})$$

$$\alpha_{DNAij} \sim N(0, \sigma^2_{DNAplant-bird})$$

Hence, we used random effects for both bird species and individual plants as well as their interaction to obtain the probability of visit from each bird species to each individual plant.

Standard deviation parameters had half-Normal priors with large standard deviations ($\sigma$ = 3) as the variation in visitation rate among bird species is usually quite large. The prior average number of visits from a given bird species on a given plant ($\mu_{birdDNA}$) was set rather low: Normal(- 6.5, 1) on logit scale, as most bird species do not visit most plants.

Finally, we calculated the posterior probability of visit from each bird species to each individual plant ($Pvisit.bird_{DNAij}$) as the product of the probability of visit for each plant at each site ($Pvisit_{DNAi}$) and the relative probability of visit for each bird species on each plant ($Pbird_{DNAij}$ in the Binomial model above).

*Estimating probability of visit from video analysis*

We used similar reasoning and models to estimate the probability of visit from each bird species to each individual plant from video records. First, we estimated the number of bird visits per hour to each individual plant *i* using a Poisson distribution:

$$Nvisit_{CAMi} \sim Poisson(\lambda_{CAMi})$$

where the log number of visits received by each individual ($\lambda_{CAMi}$) was modelled as

$$log(\lambda_{CAMi}) = \mu_{CAMvisit} + \alpha_{CAMi} + \text{offset}(log(\text{recording.time}_i))$$
$$\alpha_{CAMi} \sim N(0, \sigma^2_{CAMvisit})$$

In this equation, $\mu_{CAMvisit}$ is the average number of bird visits across all individual plants, and $\alpha_{CAMi}$ represents individual variation around that population average (i.e. a random intercept), drawn from a Normal distribution with standard deviation $\sigma_{CAMvisit}$. We included an offset term to account for different recording time among individual plants (range = c. 18 - 20 hours, Table S.B.3.2).

We used weakly informative Normal priors for all parameters: $\mu_{CAMvisit}$ had a Normal(1.4, 1) prior (in log scale), corresponding to c. 4 bird visits per hour. $\sigma_{CAMvisit}$ had a half-Normal prior with standard deviation = 1, i.e. Normal(0, 1) truncated at 0.

Once we had estimated the number of visits/h to each individual plant, we could calculate their relative probability of visit ($Pvisit_{CAMi}$) by dividing each plant's visits by the total number of bird visits to all individuals at the site.

Then, we estimated the probability that a given visit at each plant is from a given bird species ($Pbird_{CAMij}$). For that, we modelled the number of visits from each bird species *j* to each individual plant *i* following a Binomial distribution:

$$Nvisit_{CAMij} \sim Binomial(Nvisit_{CAMi}, Pbird_{CAMij})$$
$$logit(Pbird_{CAMij}) = \mu_{CAMbird} + \alpha_{CAMi} + \alpha_{CAMj} + \alpha_{CAMij}$$
$$\alpha_{CAMi} \sim N(0, \sigma^2_{CAMplant})$$
$$\alpha_{CAMj} \sim N(0, \sigma^2_{CAMbird})$$
$$\alpha_{CAMij} \sim N(0, \sigma^2_{CAMplant-bird})$$

As for barcoding data, we used random effects for both bird species and individual plants as well as their interaction to obtain the probability of visit from each bird species to each individual plant. Standard deviation parameters had half-Normal priors with large standard deviations ($\sigma$ = 3), and the prior average number of visits from a given bird species on a given plant ($\mu_{CAMbird}$) had Normal(-6.5, 1) prior on logit scale.

Finally, we calculated the posterior probability of visit from each bird species to each individual plant ($Pvisit.bird_{CAMij}$) as the product of the probability of bird visit for each plant at each site ($Pvisit_{CAMi}$) and the relative probability of visit for each bird species on each plant ($Pbird_{CAMij}$ in the Binomial model above).

*Merging of visitation estimates from DNA barcoding and video monitoring*

The parallel analyses of visitation rates from both videos and DNA barcoding data produced compatible pairwise probabilities of visit ($Pvisit.bird_{ij}$) for each plant-bird species pair at El Puntal site (Fig. S.E.1.1). We averaged the posterior distributions of both probabilities to obtain the consensus probability of visit arising from the combination of both data sources (barcoding and videos; Fig. S.E.1.2). This estimate could be interpreted as the consensus probability that a given bird visit at the site involves a particular bird species and individual lentisc plant. For most bird species, both methods produced quite similar probabilities of visit, and the consensus probability only reinforced those estimates. When DNA barcoding and videos suggested different probabilities of visit for some plant-bird species pair, the spread of each posterior distribution offered a natural weighting so that more uncertain estimates (from whichever method) had less influence on the final consensus probability.

At Las Madroñas site, where video recordings were not available, the pairwise probabilities of visit were estimated based on barcoding data only (Fig. S.E.1.3).

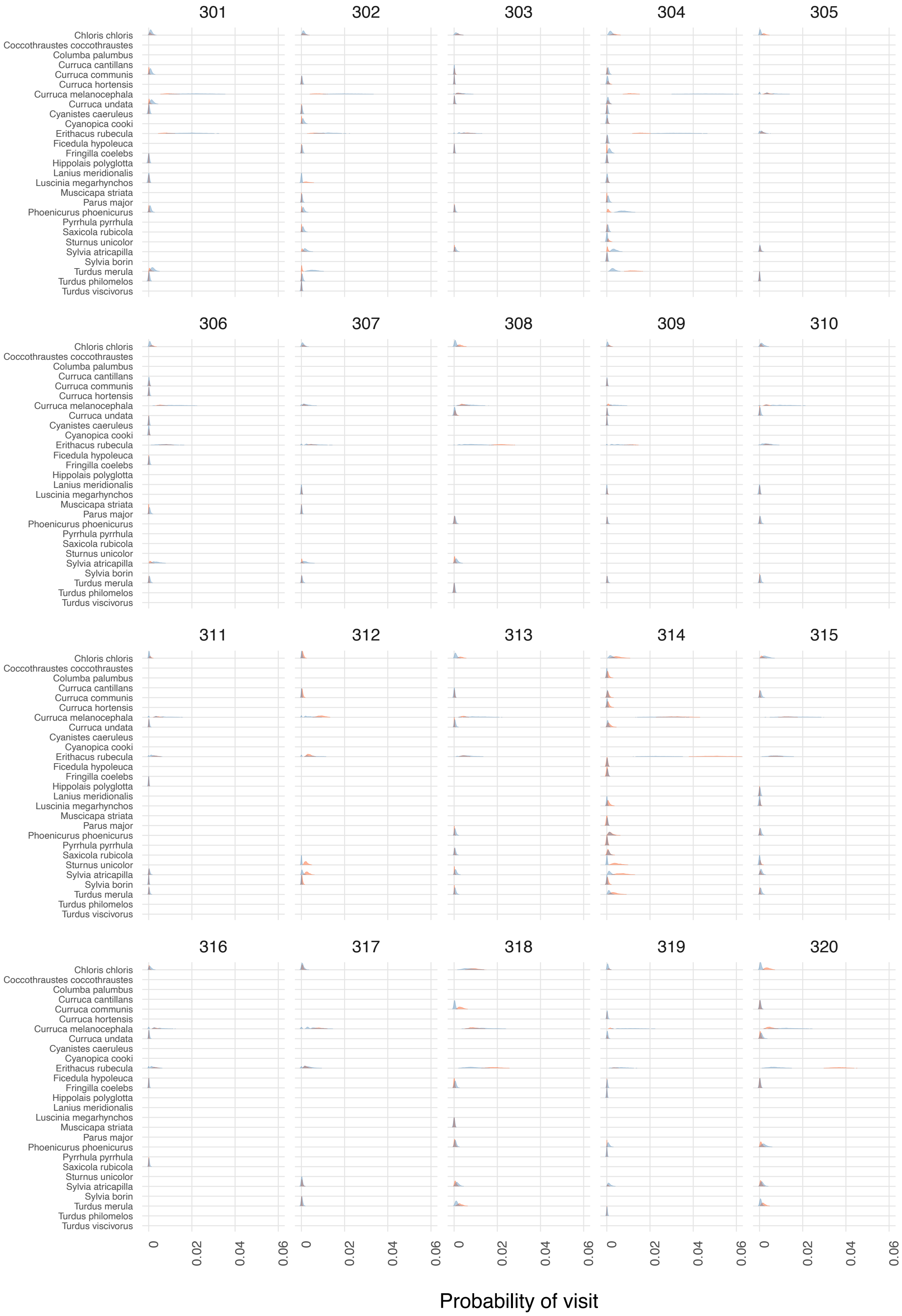
Chloris chloris
Coccothraustes coccothraustes
Columba palumbus
Curruca cantillans
Curruca communis
Curruca hortensis
Curruca melanocephala
Curruca undata
Cyanistes caeruleus
Cyanopica cooki
Erithacus rubecula
Ficedula hypoleuca
Fringilla coelebs
Hippolais polyglotta
Lanius meridionalis
Luscinia megarhynchos
Muscicapa striata
Parus major
Phoenicurus phoenicurus
Pyrrhula pyrrhula
Saxicola rubicola
Sturnus unicolor
Sylvia atricapilla
Sylvia borin
Turdus merula
Turdus philomelos
Turdus viscivorus
0
0.02
0.04
0.06
Probability of visit

continued

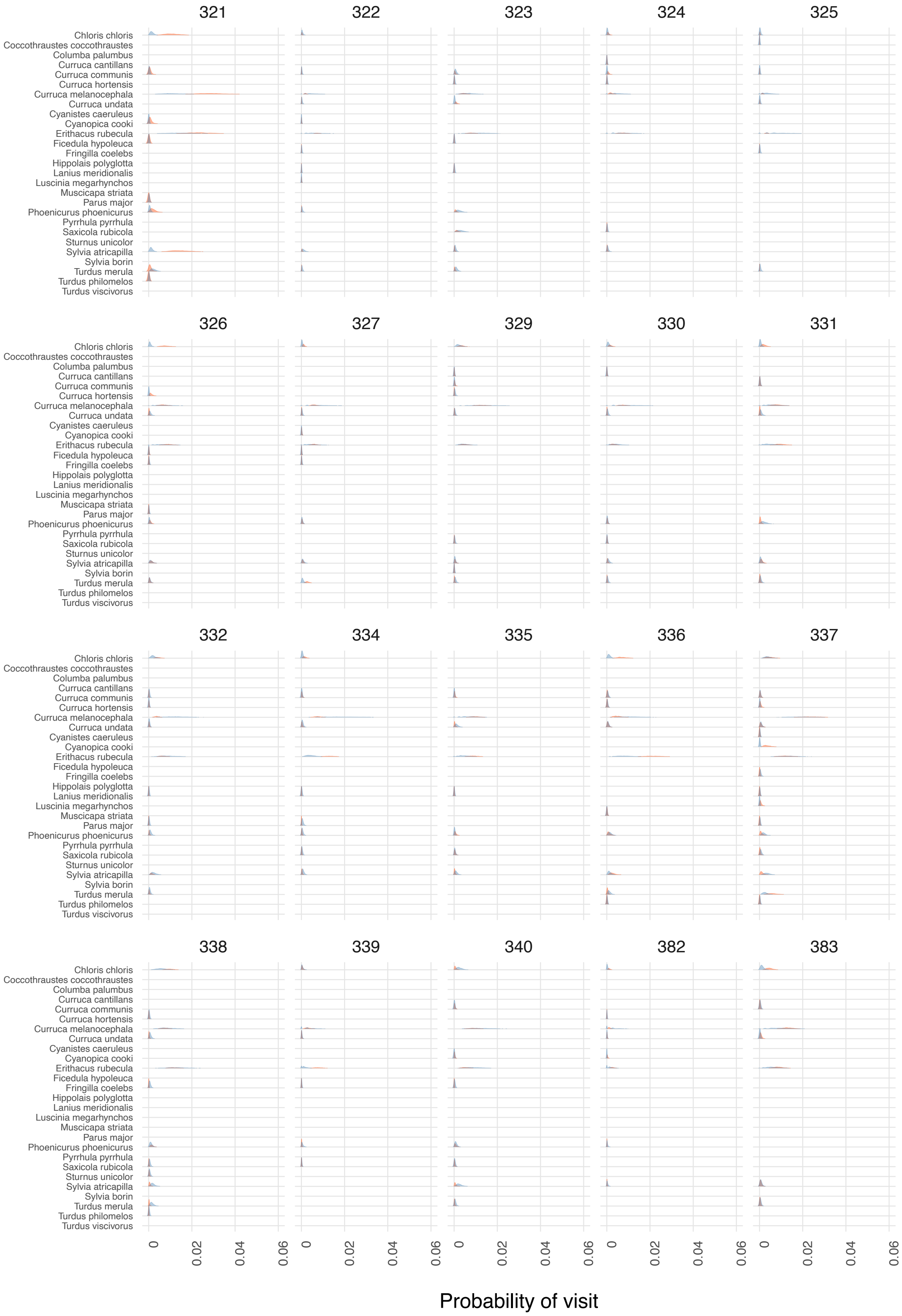

**Figure S.E.1.1.** Estimated probability of visit from each bird species to each individual plant at El Puntal site. Posterior distributions in red and blue colours represent estimates arising from DNA barcoding and video cameras, respectively. Panel numbers represent different plant IDs.

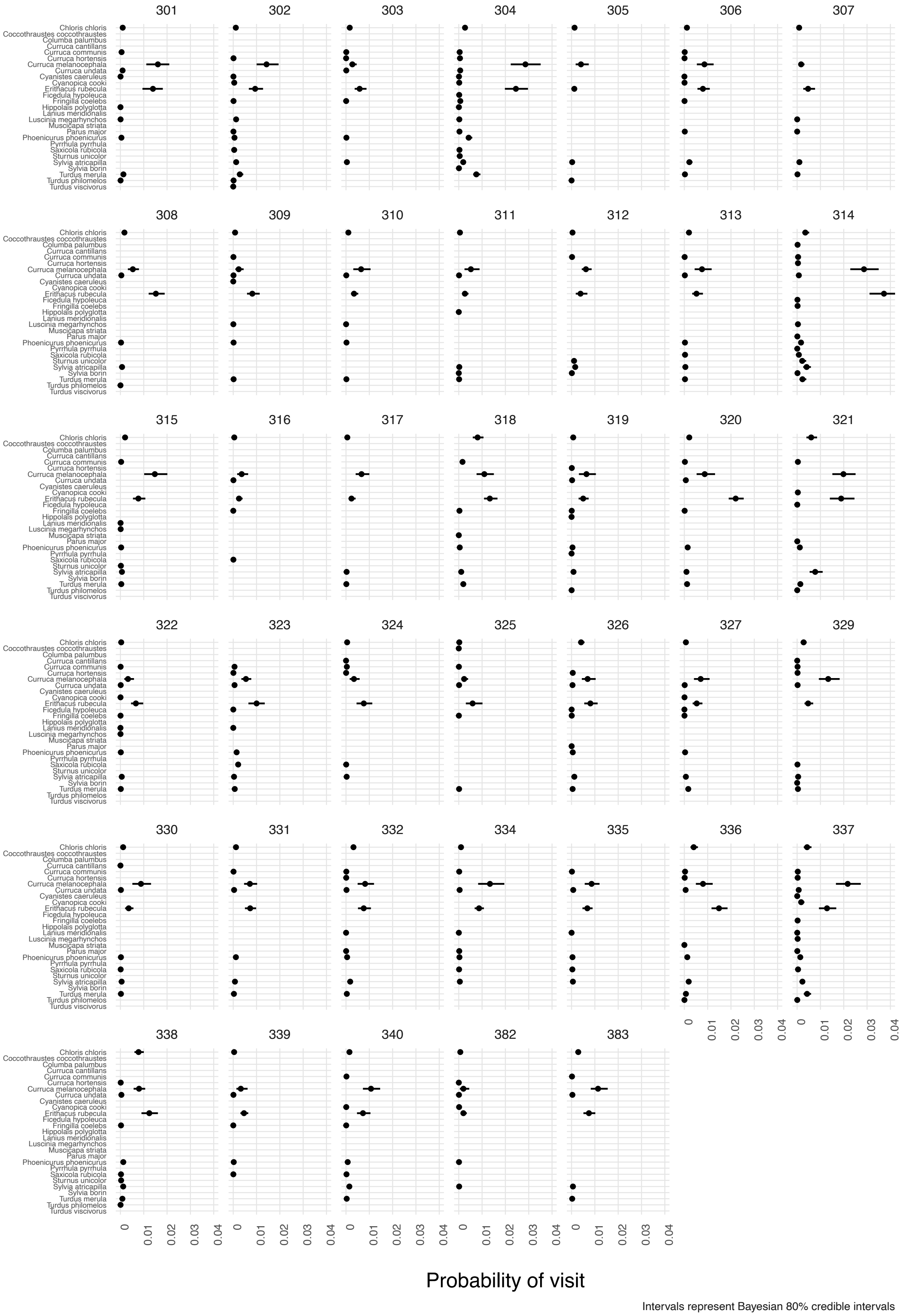

**Figure S.E.1.2.** Consensus probability of visit from each bird species to each individual plant at El Puntal site, obtained by averaging posterior distributions from DNA barcoding and video cameras.

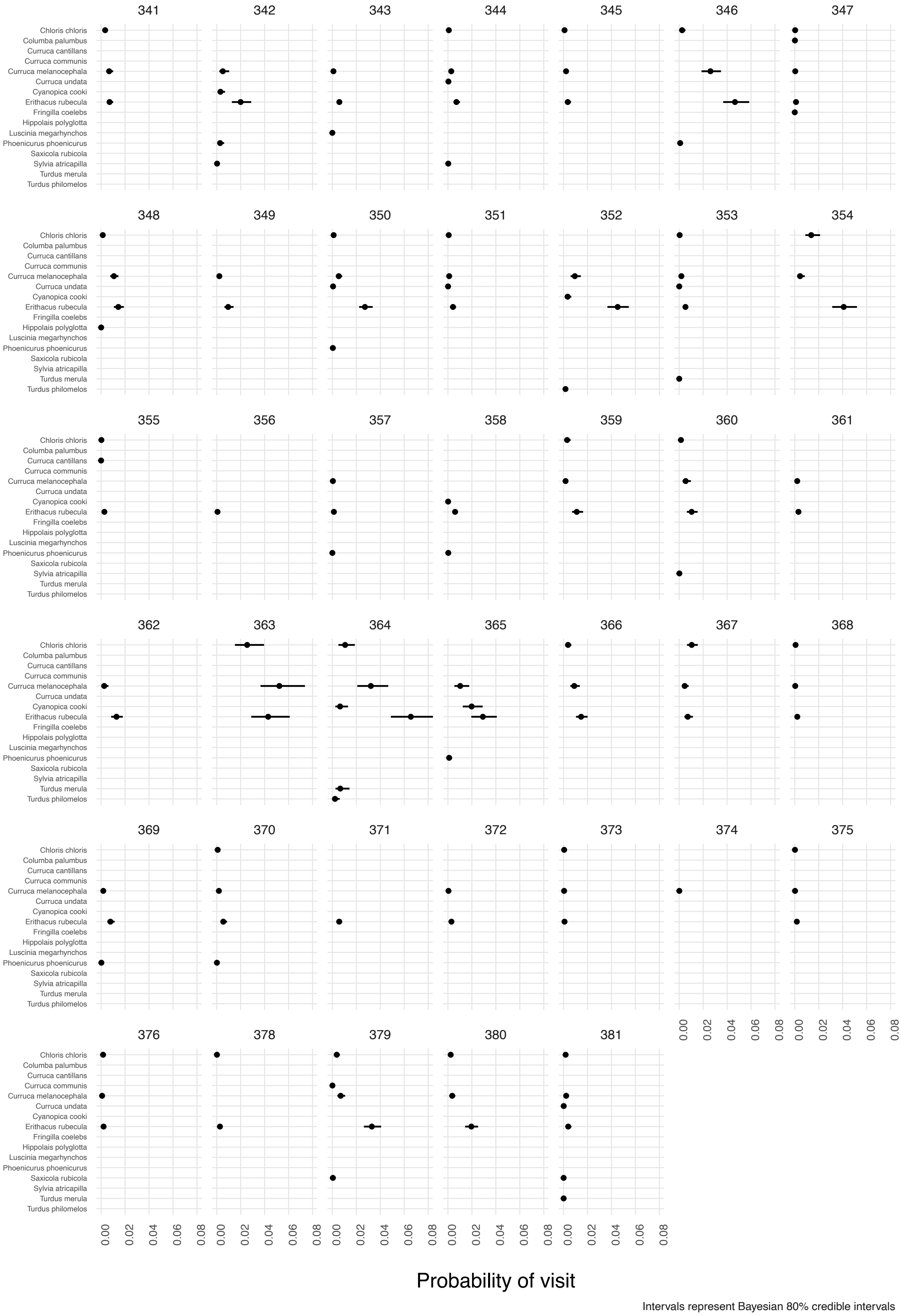

**Figure S.E.1.3.** Posterior probability of visit from each bird species to each individual plant at Las Madroñas site, estimated from DNA barcoding of bird droppings beneath each plant.

**Probability of fruit/seed consumption per visit**

To estimate the proportion of bird visits involving fruit or seed consumption we used information on feeding bouts obtained from video recordings (Fig. S.E.1.4). For each bird species we recorded the number of visits involving feeding and those where the bird left the plant without consuming any fruit or seed. To obtain the probability of feeding ($Pfeed_j$) for each bird species we analysed these data using a Bernoulli distribution and a random effect for bird species:

$$Feed_j \sim Bernoulli(Pfeed_j)$$

$$logit(Pfeed_j) = \mu_{feed} + \alpha_j$$

$$\alpha_j \sim N(0, \sigma^2_{feed})$$

$\mu_{feed}$ had a weak prior probability Normal(0, 2) on logit scale, corresponding to a very uncertain probability of feeding centred around 0.5. $\sigma_{feed}$ had the same prior N(0, 2) allowing for different visiting and feeding behaviours among bird species.

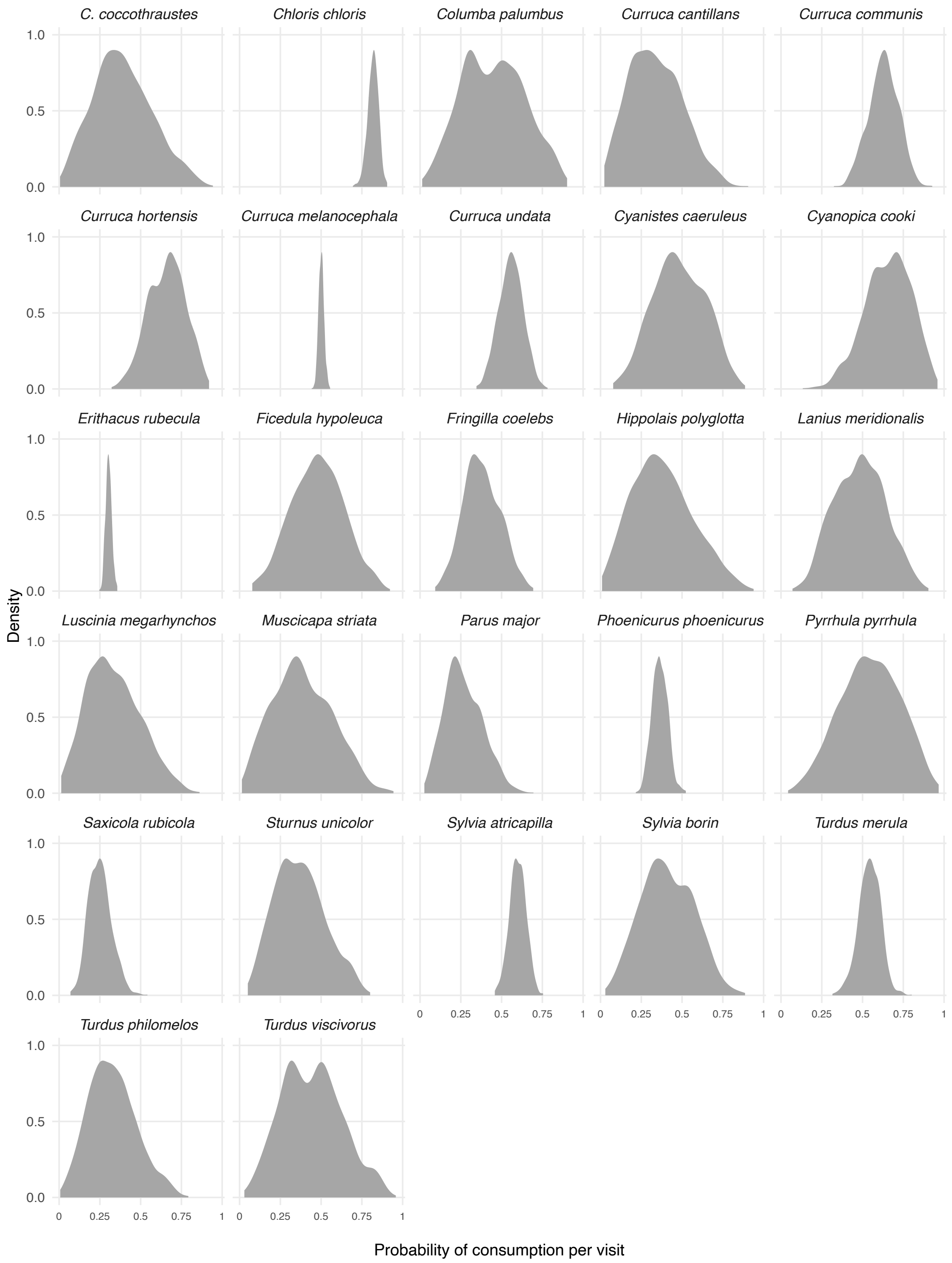

**Figure S.E.1.4.** Estimated probability for each bird species of consuming at least one fruit or seed when visiting lentisc plants.

**Number of fruits/seeds consumed per visit**

Once we had estimated the number of visits from each bird species and their probability of feeding in each visit, we estimated the number of fruits or seeds consumed in feeding visits ($Nfruit_j$; Fig. S.E.1.5) using a Negative Binomial distribution with a random effect for bird species and including body mass (from Elton Traits: Wilman *et al.* 2014) as a covariate:

$$Nfruit_j \sim NegBinomial(\lambda_j, \phi_{fruit})$$

$$log(\lambda_j) = \mu_{fruit} + \alpha_j + \beta_{BM}\, log(\text{bodymass}_j)$$

$$\alpha_j \sim N(0, \sigma^2_{fruit})$$

As 'bodymass' predictor was centred around 20 grams value, $\mu_{fruit}$ is the expected log number of fruits/seeds consumed by a bird species with 20 grams of body mass, and was assigned a Normal(0.7, 0.3) prior distribution on log scale, corresponding to an expected grand mean of 2 fruits consumed per visit. $\alpha_j$ is the bird species random effect. Its standard deviation $\sigma_{fruit}$ had a half-Normal prior with standard deviation of 0.5 units. The $\beta_{BM}$ parameter represents the expected increase in fruit consumption with increasing body mass and had a weakly informative Normal(0.5, 0.5) positive prior since the amount of fruits consumed is generally positively associated to bird size. Finally, the $\phi_{fruit}$ parameter accommodates overdispersion in the count data (Winter & Bürkner 2021) and had a Gamma(0.01, 0.01) prior distribution.

Since we modelled the probability of consumption independently, here we only used feeding observations where at least one fruit or seed was consumed. Thus, we used a lower truncation value of one. Also, since *Chloris chloris* is an eager seed predator with radically different feeding behaviour, we modelled this species independently to preserve the assumption of exchangeability of the random effects (Kéry & Schaub 2021). In this case we ran an intercept-only truncated negative binomial model with Normal(2, 0.5) prior distribution on log scale, corresponding to a mean of c. 7.4 seeds consumed per visit.

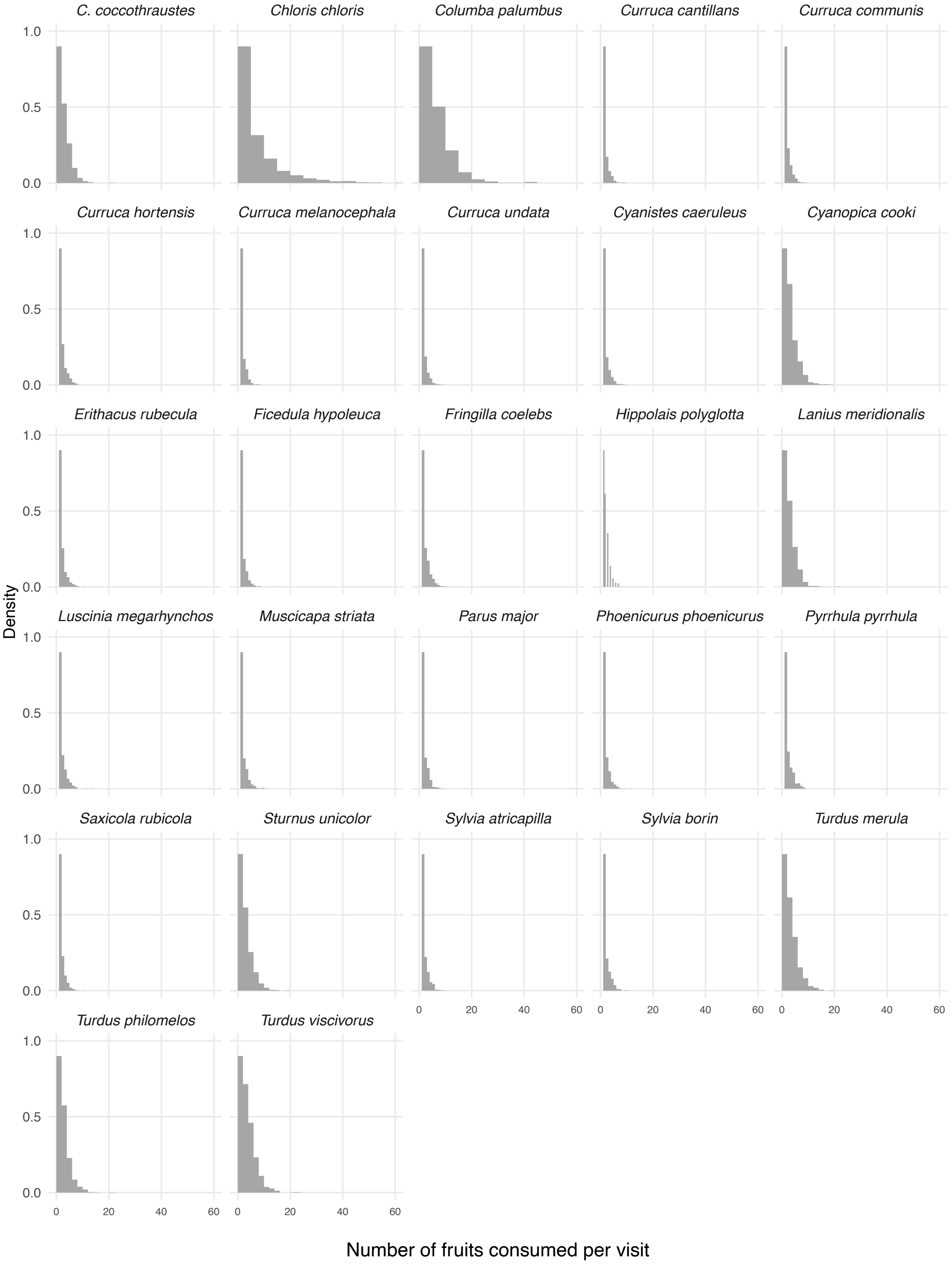

**Figure S.E.1.5.** Estimated number of fruits consumed per visit by each bird species.

**Calculating the quantity component**

To calculate the number of fruits/seeds consumed by each bird species from each plant (the quantity component QTY for both effectiveness estimates RPE and SDE), we multiplied all the posterior distributions estimated previously, namely (i) the total number of bird visits at each site *k* ($Nvisit_k$), (ii) the probability of visit from each bird species *j* to each plant *i* ($Pvisit.bird_{ij}$), (iii) the probability that a visit from a bird species *j* involves fruit or seed consumption ($Pfeed_j$), and (iv) the number of fruits/seeds consumed per feeding visit ($Nfruit_j$):

$$QTY_{ij} = Nvisit_k \times Pvisit.bird_{ij} \times Pfeed_j \times Nfruit_j$$

*E.2. Quality component of Resource Provisioning Effectiveness*

To estimate the quality of individual plants' reward we calculated the energy acquired per fruit (for pulp consumers) or seed consumed (for granivorous birds). Energy calculations were based on: seed/pulp fresh mass, percentage of water in seed/pulp and the energetic yield factors for seed and pulp dry mass.

Pulp and seed fresh mass of individual plants (Suppl. Mat. C) was converted to dry mass using information available on % water content in seed and pulp reported in nearby *P. lentiscus* populations (Jordano 1984). To estimate the energy rendered per g of dry seed mass, we used a yield factor of 28.14 kJ/g (Khiari *et al.* 2020). Pulp yield energy was estimated based on lipid, carbohydrate and protein percentages (5.5 % proteins, 58.8 % lipids and 25.8 % carbohydrates; Herrera 1987). These percentages were multiplied by standard energy conversion factors for major nutrients in fruits (14.1 kJ/g for proteins, 35.0 kJ/g for lipids and 15.1 kJ/g for carbohydrates; MacLean *et al.* 2003). The resulting energetic yield factor for pulp dry mass was 25.25 kJ/g.

Final quality was then calculated as the product of seed/pulp fresh mass (g), non-water % in seed/pulp, and seed/pulp yield energy factor (kJ/g). Variations in seed and fruit quality between individual plants were therefore based on differences between the fresh mass of pulp and seed.

*E.3. Quality component of Seed Dispersal Effectiveness*

#### Probability of escaping granivorous birds predation

For granivorous birds (*Chloris chloris, Pyrrhula pyrrhula, Fringilla coelebs and Coccothraustes coccothraustes*) we estimated the probability of seeds escaping predation as they are dropped during handling. This is a very rare event, but it happens occasionally. To estimate its frequency we counted the number of intact and destroyed seeds collected in seed traps beneath lentisc plants, and fitted a hierarchical Binomial regression:

$$Ndropped_i \sim Binomial(Nseed_i, Pdrop_i)$$

$$logit(Pdrop_i) = \mu_{drop} + \alpha_i$$

$$\alpha_i \sim N(0, \sigma^2_{drop})$$

where the proportion of intact (dropped) seeds beneath each mother plant, or probability of escaping predation ($Pdrop_i$), is modelled as a random effect with mean $\mu_{drop}$ with a weak Normal(-6.9, 2) prior distribution (corresponding to one seed per thousand escaping predation on average) and standard deviation $\sigma_{escape}$ with a half-Normal(0, 1) prior. This analysis reported a posterior probability of escaping predation of 0.0014 ± 0.0005 (mean ± SE).

#### Probability of dispersal to different microhabitats

For each bird species we estimated the probability of dispersing seeds to each of the five microhabitats defined (PL: under *Pistacia lentiscus* plants, FR: under other fleshy fruited species, NF: under non-fleshy fruited species, OA: open ground areas, P: under pine trees). For that we used two steps: first we modelled the total number of seeds arriving to each microhabitat, and then we identified the proportion of seeds brought by each bird species using the identifications obtained through DNA barcoding of bird droppings.

To estimate the seed rain density in each microhabitat we modelled the total number of seeds arriving per $m^2$ using a Negative Binomial distribution:

$$Nseed_s \sim NegBinomial(\eta_s, \phi_s)$$

$$log(\eta_s) = \mu_{FR} + \mu_m + \text{offset}(log(\text{sampling.area}_s))$$

Here, $Nseed_s$ is the total number of seeds collected at each seed trap or transect, $\mu_{FR}$ is the average number of seeds arriving per $m^2$ in the FR microhabitat (taken as intercept), $\mu_m$ is the average difference between each microhabitat and FR, and $\phi_s$ accommodates overdispersion in the count data. $\mu_{FR}$ had a weakly informative Normal(3, 2) prior centred around 20 seeds/m$^2$, and $\mu_m$ had a Normal(0, 2) prior, allowing for large differences in seed rain density among microhabitats. $\phi_s$ had a Gamma(0.01, 0.01) prior. We used an offset to account for different sampling area across microhabitats.

To estimate the proportion of seeds contributed by each bird species *j* to each sampling station *s* in each microhabitat *m*, we used a hierarchical Binomial model:

$$Nseed_{sj} \sim Binomial(Nseed_s, Pseed.bird_{mj})$$

$$logit(Pseed.bird_{mj}) = \beta_{FRj} + \beta_{NFj} + \beta_{OAj} + \beta_{PLj} + \beta_{PPj}$$

$$\begin{pmatrix} \beta_{FRj} \\ \beta_{NFj} \\ \beta_{OAj} \\ \beta_{PLj} \\ \beta_{PPj} \end{pmatrix} \sim N\left(\begin{pmatrix} \mu_{\beta_{FRj}} \\ \mu_{\beta_{NFj}} \\ \mu_{\beta_{OAj}} \\ \mu_{\beta_{PLj}} \\ \mu_{\beta_{PPj}} \end{pmatrix}, \begin{pmatrix} \sigma^2_{\beta_{FRj}} & \rho_{\beta_{FRj}\beta_{NFj}} & \rho_{\beta_{FRj}\beta_{OAj}} & \rho_{\beta_{FRj}\beta_{PLj}} & \rho_{\beta_{FRj}\beta_{PPj}} \\ \rho_{\beta_{NFj}\beta_{FRj}} & \sigma^2_{\beta_{NFj}} & \rho_{\beta_{NFj}\beta_{OAj}} & \rho_{\beta_{NFj}\beta_{PLj}} & \rho_{\beta_{NFj}\beta_{PPj}} \\ \rho_{\beta_{OAj}\beta_{FRj}} & \rho_{\beta_{OAj}\beta_{NFj}} & \sigma^2_{\beta_{OAj}} & \rho_{\beta_{OAj}\beta_{PLj}} & \rho_{\beta_{OAj}\beta_{PPj}} \\ \rho_{\beta_{PLj}\beta_{FRj}} & \rho_{\beta_{PLj}\beta_{NFj}} & \rho_{\beta_{PLj}\beta_{OAj}} & \sigma^2_{\beta_{PLj}} & \rho_{\beta_{PLj}\beta_{PPj}} \\ \rho_{\beta_{PPj}\beta_{FRj}} & \rho_{\beta_{PPj}\beta_{NFj}} & \rho_{\beta_{PPj}\beta_{OAj}} & \rho_{\beta_{PPj}\beta_{PLj}} & \sigma^2_{\beta_{PPj}} \end{pmatrix}\right)$$

The probability that a seed arriving at a given microhabitat is brought by bird species *j* was modelled as a random effect where parameters ($\beta_{FRj}$, $\beta_{NFj}$, $\beta_{OAj}$, $\beta_{PLj}$, $\beta_{PPj}$) are drawn from a multivariate Normal distribution. $\mu_{\beta FRj}$ had a weak Normal(-3.3, 1) prior (assuming equal prior probability among bird species as the inverse logit of -3.3 ≈ 1/27 bird species), the $\sigma$ parameters had half-Normal(0, 2) priors, and the correlation matrix among $\beta$ parameters had LKJ(2) prior distribution.

Then, to estimate the number of seeds dispersed to each microhabitat by each bird species we multiplied the posterior from the first model (total seed rain per microhabitat) with the probability that seeds are brought by each bird species (model above). For each bird species, the relative probability of dispersing seeds to each microhabitat (Fig. S.E.3.1) can finally be calculated as the ratio of the number of seeds dispersed to each microhabitat by the total number of seeds dispersed.

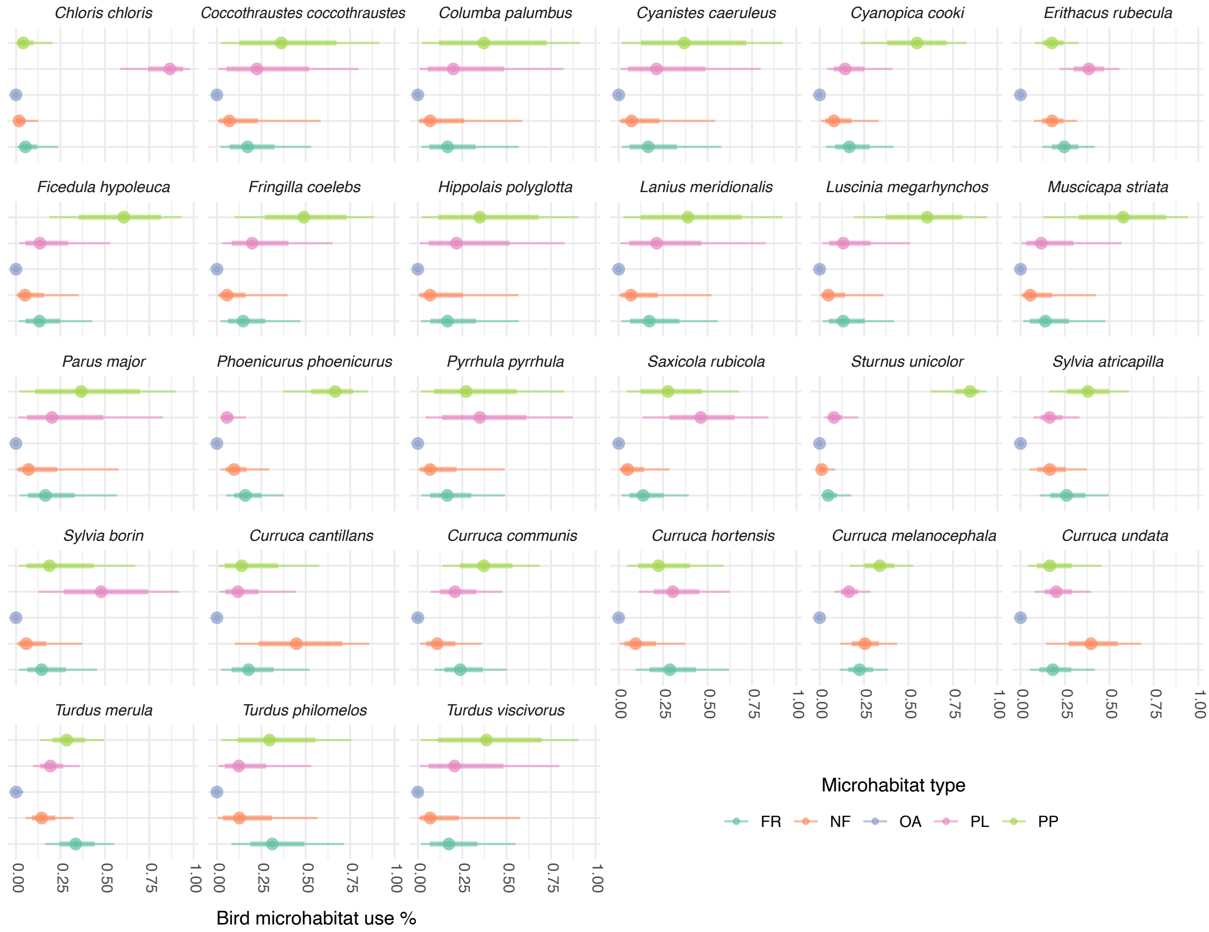

**Figure S.E.3.1.** Posterior probabilities of dispersal to each microhabitat by each of the 27 bird species consuming *Pistacia lentiscus* fruits or seeds (FR: under fleshy fruited species (non–*Pistacia*), NF: under non-fleshy fruited species, OA: in open ground areas, PL: under *Pistacia lentiscus* stands, PP: under pine trees). Dots denote median probability, and thick and thin bars represent 66% and 95% credible intervals, respectively.

### Probability of escaping post-dispersal predation

Rodent predation was quite fast and severe. Two days after installing the experiment, half of the experimental units had received full or partial predation. Within the first month more than 90% of the open units had experienced predation. At the same time, our seedling emergence experiments showed that seedlings started emerging on the 28th day after sowing, which agrees with published evidence of early germination and emergence in the species (García-Fayos & Verdú 1998; Del Campo *et al.* 2014). Therefore we considered the first month after dispersal as the critical period for seeds to be preyed upon before germination.

Our mesh-protected experimental units failed to prevent rodent predation on several occasions, hence we discarded using the data from these controls for the analysis of predation escape. In those control units that effectively repelled rodents, seeds typically remained intact for a long time, suggesting that most seed predation is actually done by rodents.

To estimate the probability of surviving post-dispersal predation in each microhabitat (Fig. S.E.3.2), we thus counted the number of intact (not predated) seeds within the first month after installing the experiment. We used a Binomial distribution:

$$Nintact_u \sim Binomial(10, Pescape_u)$$

$$logit(Pescape_u) = \mu_{escapeFR} + \beta_{escapeNF} + \beta_{escapeOA} + \beta_{escapePL} + \beta_{escapePP} + \alpha_u$$

$$\alpha_u \sim N(0, \sigma^2_{escape})$$

$\mu_{escapeFR}$ represents the probability of escaping predation in the FR (fleshy-fruited) microhabitat, taken as intercept, and was given a weak Normal(-1, 2) prior implying relatively high predation rates (as rodent predation is often higher under dense vegetation; Fedriani & Manzaneda 2005). The $\beta$ parameters thus represent the differences in predation rates in the other microhabitats (compared to FR), and were given broad Normal(0, 2) prior distributions allowing for large differences between microhabitats. Finally, we included an observation-level random effect ($\alpha_u$) to account for potentially overdispersed predation rates among experimental units, with a half-Normal(0, 3) prior for $\sigma_{escape}$.

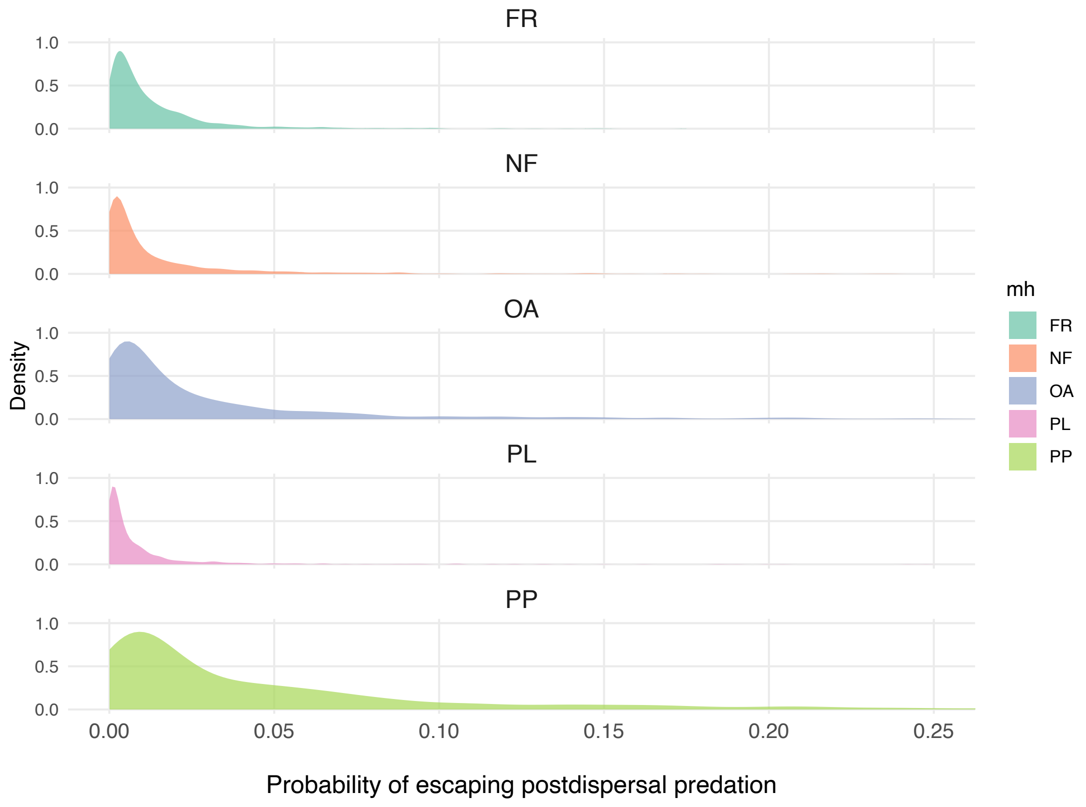

**Figure S.E.3.2.** Posterior probability of escaping post-dispersal seed predation in each microhabitat.

#### Probability of seedling emergence and survival

We estimated the probability of emergence and seedling survival through the first summer (up to mid October) using data from two seasons (2018-19 and 2019-20) (Fig.S.E.3.3):

$$Survival_s \sim Bern(Psurv_s)$$

$$logit(Psurv_s) = \mu_{survFR} + \beta_{survNF} + \beta_{survOA} + \beta_{survPL} + \beta_{survPP} + \beta_{2020} + \beta_{survNF2020} + \beta_{survOA2020} + \beta_{survPL2020} + \beta_{survPP2020} + \alpha_e$$

$$\alpha_e \sim N(0, \sigma^2_{surv})$$

Seedling emergence and survival was modelled as a Bernoulli process, with probability depending on microhabitat and season. $\mu_{survFR}$, the intercept parameter, represents the probability of survival on the FR (fleshy-fruited) microhabitat in the first season, and was given a Normal(-6.9, 2) prior (logit scale), corresponding to 0.1% survival (i.e. only 1 in 1000 dispersed seeds would produce a seedling still alive after their first summer). The $\beta$ parameters accommodate differences in survival probability among microhabitats and seasons, and had Normal(0, 2) priors. Finally, there was a random effect ($\alpha_e$) to account for replicated measurements within sowing units (each experimental unit had 16 sown seeds). $\alpha_e$ was drawn from a Normal distribution with standard deviation $\sigma_{surv}$ having half-Normal(0, 1) prior.

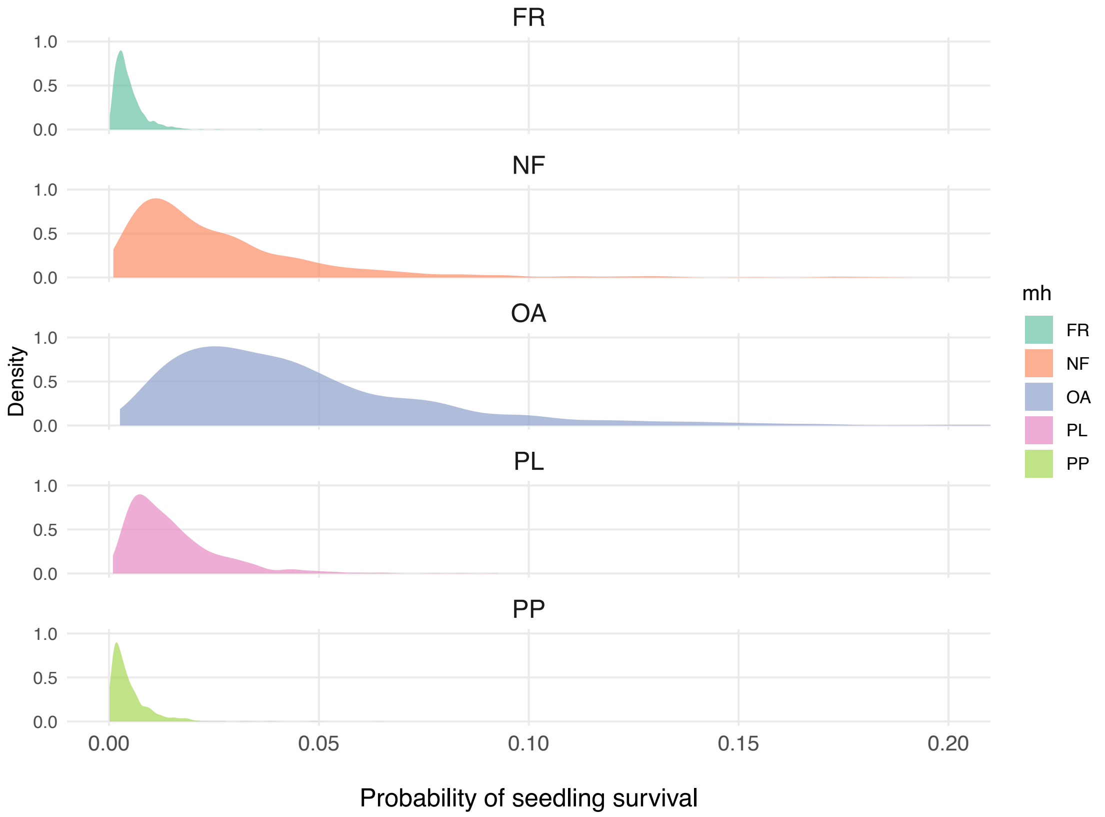

**Figure S.E.3.3.** Posterior probability of seedling emergence and early survival in each microhabitat, combining results from both 2018-19 and 2019-20 seasons.

#### Calculating the quality component of SDE

The quality component of seed dispersal effectiveness (SDE) estimates the probability of a seed dispersed by a given bird species to turn into a seedling surviving its first summer (Fig. S.E.3.4). This probability can be calculated as the product of the posterior probabilities obtained above:

- Probability of escaping predation from granivorous birds (*Pdrop*)
- Probability, for each bird species, of dispersing seeds towards each microhabitat (*Pseed.bird*)
- Probability of escaping post-dispersal (rodent) predation in each microhabitat (*Pescape*)
- Probability of emergence and early seedling survival in each microhabitat (*Psurv*)

Note these estimates of SDE quality represent an upper bound of seedling recruitment per dispersed seed since we do not account for the viability of dispersed seeds (González-Varo *et al.* 2019).

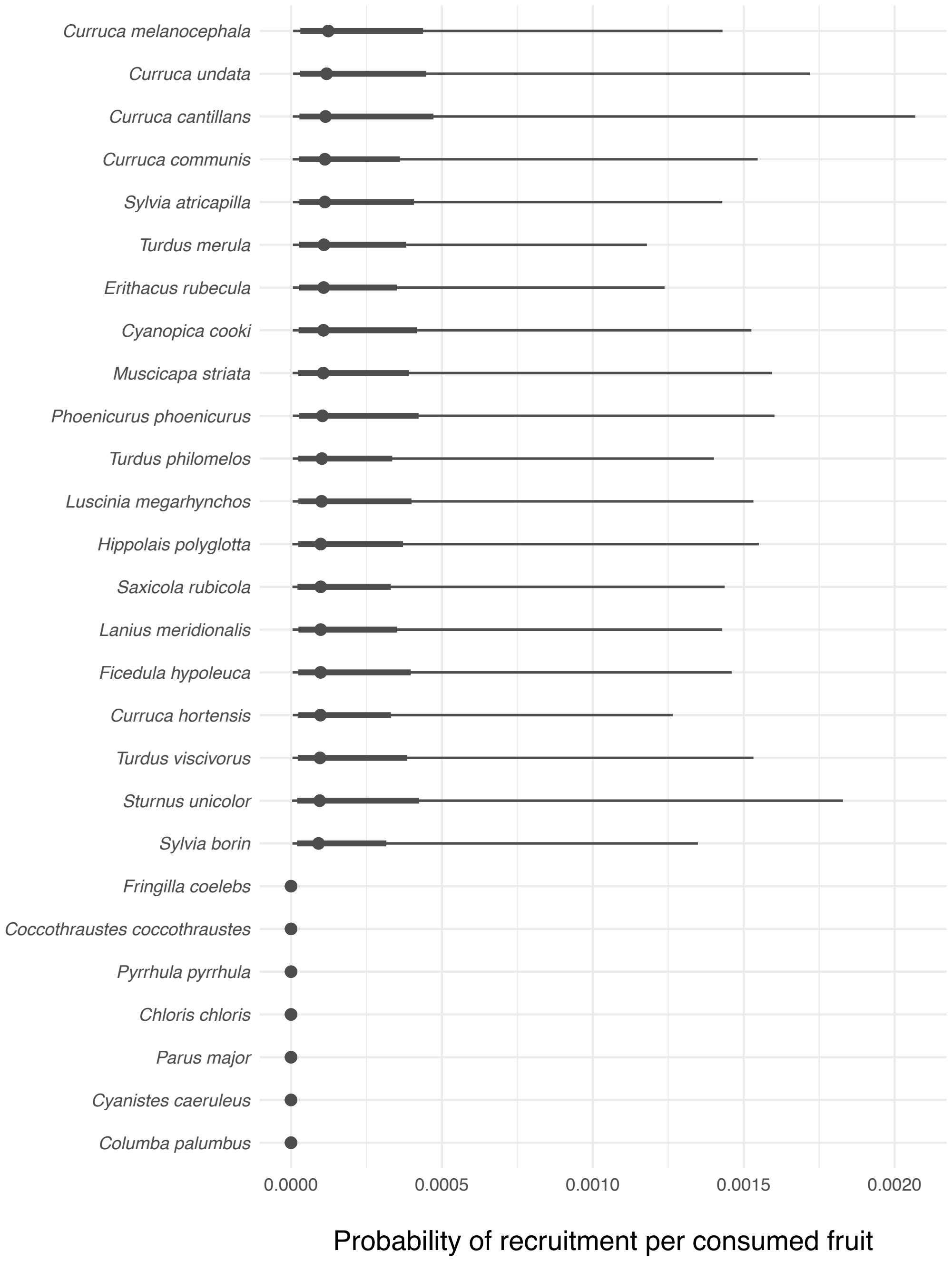

**Figure S.E.3.4.** The quality component of seed dispersal effectiveness, represented as the probability of seedling recruitment per consumed fruit or seed for each bird species. Bird species appear sorted by decreasing median probability (represented by dots). Intervals represent Bayesian 66% and 95% credible intervals (thick and thin lines, respectively).

*E.4. Complete SDE landscape (including non-legitimate dispersers)*

Seed predators and pulp peckers were removed from the Seed Dispersal Effectiveness (SDE) landscape shown in the main text to facilitate visualisation of the quality component and the differences between legitimate dispersers. The following figure S.E.4.1 shows the complete SDE landscape incorporating the *non-legitimate dispersers* (*Chloris chloris*, *Fringilla coelebs, Pyrrhula pyrrhula, Coccothraustes coccothraustes*, *Columba palumbus, Parus major* and *Cyanistes caeruleus*).

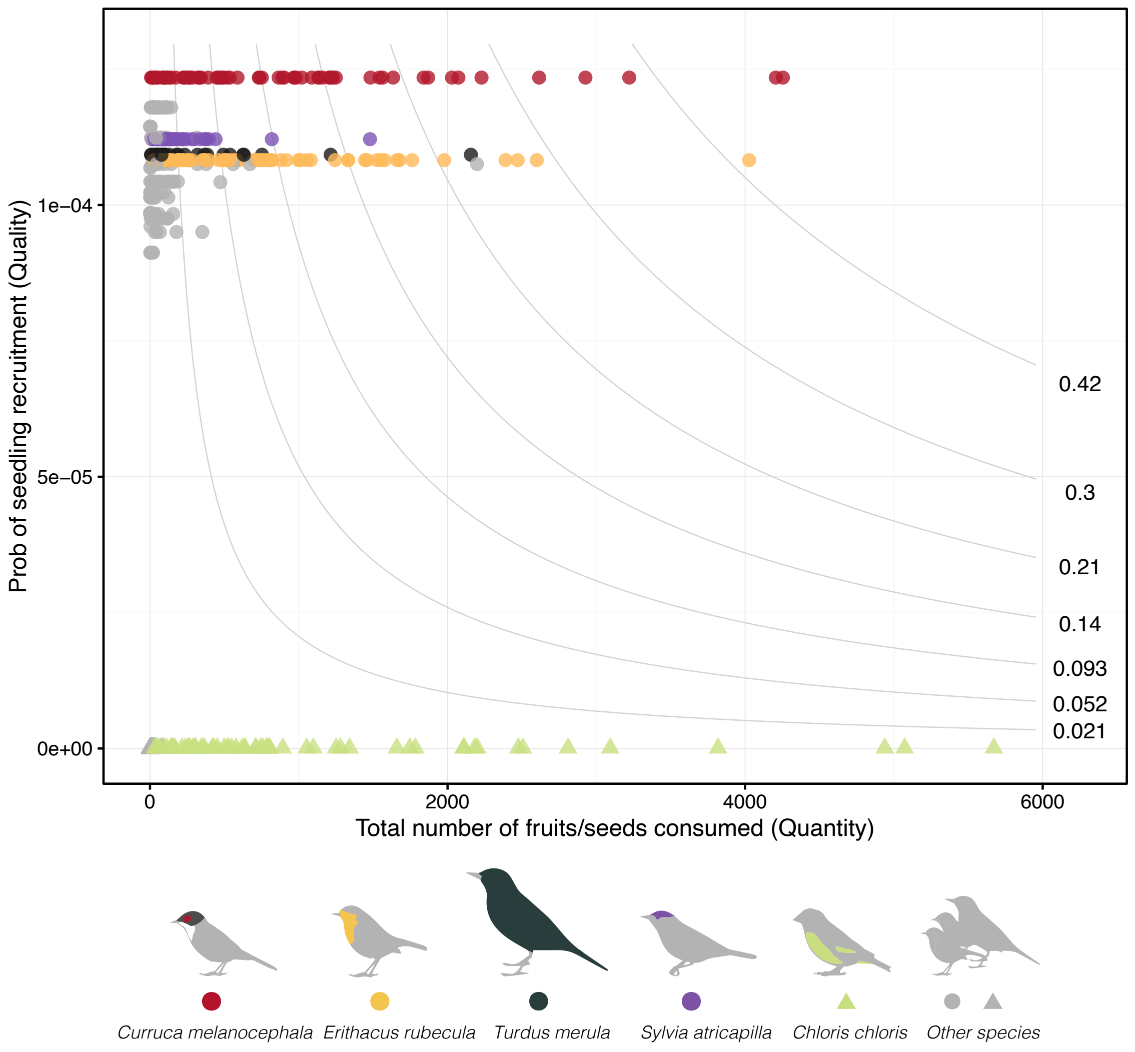

**Figure S.E.4.1.** Seed dispersal effectiveness landscape (SDE) for individual *Pistacia lentiscus* plants. Each point represents an individual plant pairwise interaction with a given avian frugivore species represented in different colours. The horizontal axis depicts the total number of fruits (or seeds, in the case of the granivorous species) consumed by each bird species in each individual plant and the vertical axis represents the posterior median probability of recruiting a seedling from a fruit ingested by each bird species. The product of the horizontal (Quantity) and vertical (Quality) axis gives the total number of plant recruits for each bird-plant pairwise interaction. Different combinations of quantity and quality can result in equal effectiveness values, as shown by the SDE isolines.

*E.5. Variance partitioning of effectiveness components*

To estimate the relative importance of each component (i.e., quantity and quality) on the total effectiveness, we adjusted separate models of effectiveness as a function of each component (all variables were log-transformed). For Seed Dispersal Effectiveness we only considered interactions with legitimate dispersers. The coefficient of determination ($R^2$) of each model represented the partitioned variance of the total effectiveness. $R^2$ values were normalised to sum up to 100%.

### F. Reciprocity and Asymmetry calculations

#### *F.1. Analysis of reciprocity*

Reciprocity between the reward of individual plants and frugivorous birds was estimated using Pearson correlation coefficients between the log-transformed RPE and SDE values. We aggregated the total rewards offered and received by each individual plant (i.e. adding up the rewards across all bird species interacting with each plant), using the 1000 posterior distribution samples (Fig. S.F.1.1). A high positive correlation between RPE and SDE would indicate high reciprocity: individual plants contributing high resource provisioning (RPE) obtain in turn high dispersal effectiveness (SDE) from their assemblage of frugivores.

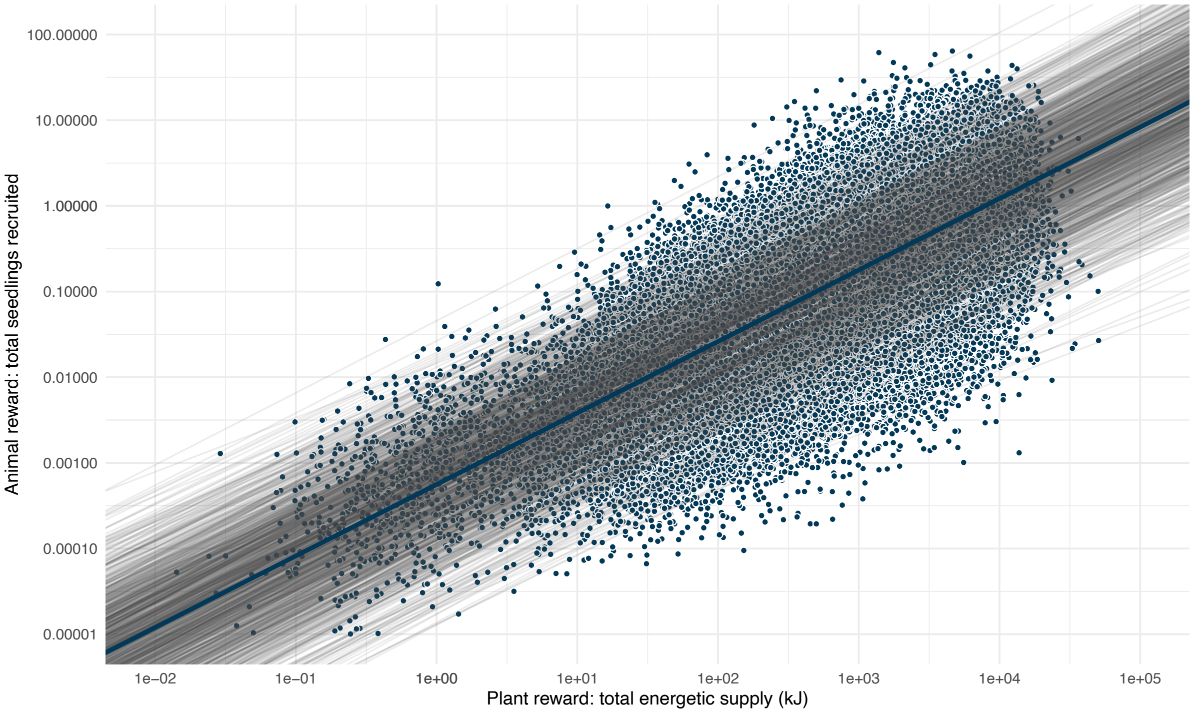

**Figure S.F.1.1** Relationship between the total energetic supply provided by individual plants (aggregating all its consumer bird species) and the number of seedlings recruited by each plant (n = 79). Each point represents one of the 1000 posterior distribution probabilities estimated per plant. Grey shaded lines represent the linear trend for each of the 1000 posteriors, and the thicker dark line represents the mean linear trend. Note both axes are in logarithmic scale.

*F.2. Dependence and asymmetry calculations*

We calculated mutual dependence (*d*) for each pairwise interaction, so that two dependence values were obtained: $d_{P_i \rightarrow A_j}$ measures the proportion of seeds dispersed (SDE) that plant *i* receives from animal species *j* relative to all the seeds dispersed for that plant. In turn, $d_{A_j \rightarrow P_i}$ measures the proportion of energy acquired (RPE) that animal species j receives from individual *P. lentiscus* plant *i,* relative to all the energy acquired by that animal (eq. 1). The sum of the dependencies of a given species/individual on all its partners must equal 1.

eq. 1a: $d_{P_i \rightarrow A_j} = \dfrac{SDE_{ij}}{\sum_{A=1}^{n} SDE_i}$, for the dependence of *P. lentiscus* plant *i* on animal species *j*;

and

eq. 1b: $d_{A_j \rightarrow P_i} = \dfrac{RPE_{ji}}{\sum_{P=1}^{m} RPE_j}$, for the dependence of animal species *j* on plant *i,*

where *d* is the dependence of plant *i* on animal species *j*, or vice versa; $SDE_{ij}$ is the estimated number of seedlings recruited by plant *i* via frugivore species *j*; $RPE_{ji}$ is the amount of kilojoules plant *i* reported to frugivore species *j*; and *n* and *m* represent the total number of animal species and individual plants, respectively.

Interaction asymmetry (AS) is defined as the difference of animal $d_{ji}$ and plant $d_{ij}$ dependencies divided by the maximum dependence value of these two (Bascompte *et al.* 2006; Vázquez *et al.* 2007)

eq. 2: $AS_{P_i A_j} = \dfrac{d_{P_i \rightarrow A_j} - d_{A_j \rightarrow P_i}}{max(d)}$

AS values can range from -1 to 1, where 0 indicates total symmetry (i.e. both partners depend on each other with the same intensity), values approaching +1 indicate that the plant is more dependent on the animal than *vice versa*, and negative values indicate that the animal is more dependent on the plant than the plant on the animal.

To account for the potential effect of matrix size (i.e., variation in the number of individual plants surveyed) on asymmetry values, we carried out simulations with adjacency matrices including variable numbers of plants (see Suppl. Mat. H). We did not find evidence for asymmetry values being significantly biassed by changes in matrix size. In addition, we compared the observed asymmetry distribution with two different null models (see Suppl. Mat. H). We observed that the highly skewed asymmetry distribution pattern did not differ when animals and plants were allowed to interact randomly following Patefield and Vázquez null models (Patefield 1981; Vázquez et al. 2007; Dormann et al. 2009).

### G. Effects on consumption (quantity component)

#### *G.1. Proportion of plants' crop consumed*

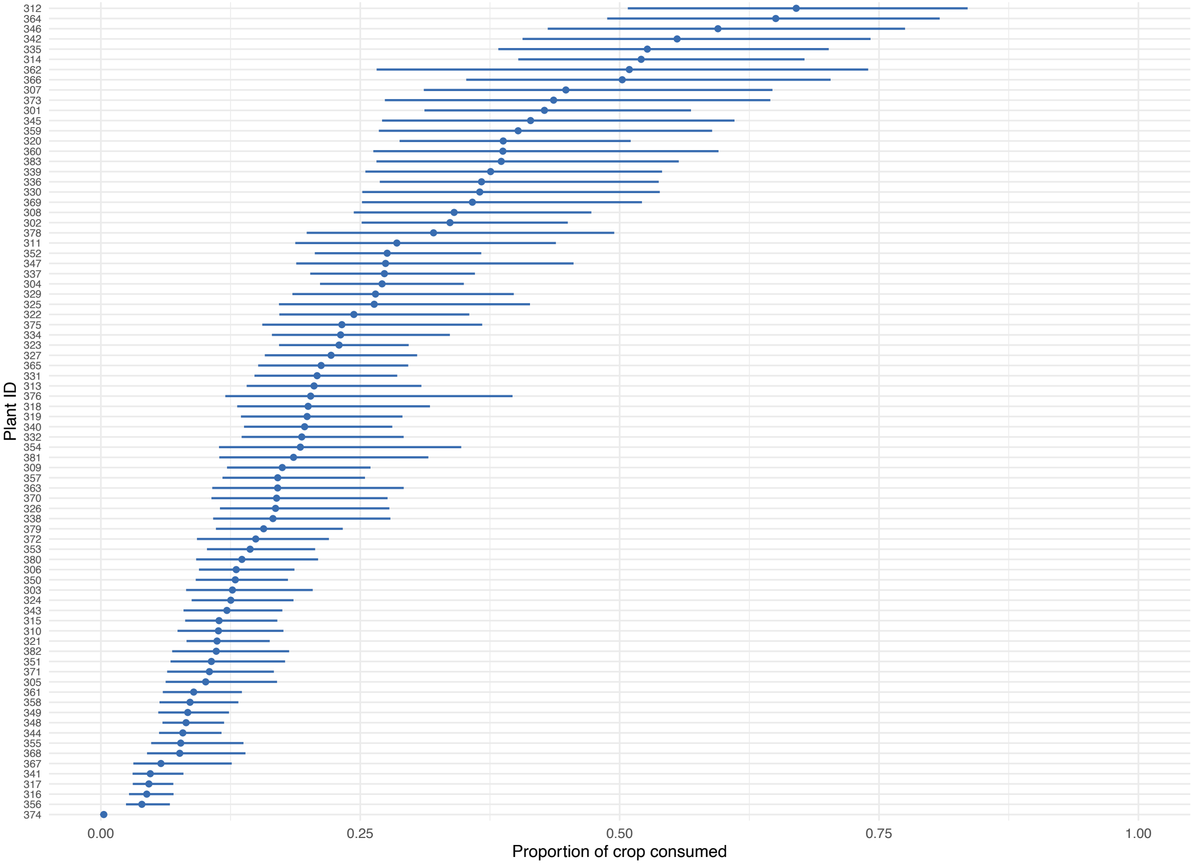

**Figure S.G.1.1.** Proportion of the initial fruit crop size estimated to be consumed by birds for each individual plant. Error bars denote the 50% credible interval.

*G.2. Birds consumption of available energy*

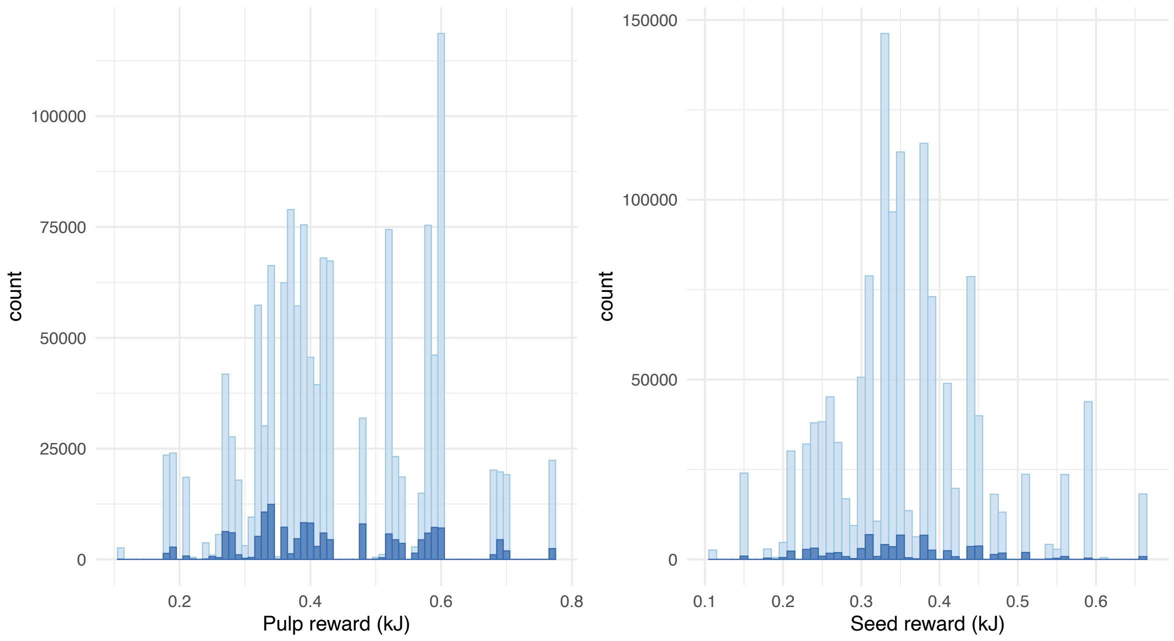

**Figure S.G.2.1.** Frequency distributions of available “pulp reward” (left) and “seed reward” (right) in kilojoules (kJ) (i.e., counts of individual fruits/seeds available at the start of the fruiting season with a given energy content per fruit or seed; light blue bars) and the estimated number of fruits/seeds consumed by birds (dark blue bars; i.e., counts of individual fruits/seeds consumed as a function of their energy content). Pulp “reward” illustrates the potential energy gain for frugivores consuming the fruits and regurgitating or defecating the seed (e.g., legitimate seed dispersers and/or pulp consumers) and seed “reward” indicates the potential energy gain for avian seed predators that discard fruit pulp and consume the seed.

*G.3. Predictors of fruit consumption intensity from individual plants*

To test the effect of different predictors (energetic pulp reward, plant canopy area and crop size) on fruit consumption from individual plants, we used a generalised linear model with a negative binomial distribution fitted with glmmTMB R-package (Brooks *et al*. 2017). All continuous predictors were log-transformed. See table S.G.3.1 with model results.

**Table S.G.3.1.** Summary statistics of the generalised linear model performed to test the effects of plant traits on fruit consumption by legitimate seed dispersers. We used a negative binomial distribution with a log link.

| **Predictors** | **Estimate ± SE** | ***p*** |
|---|---|---|
| Intercept | 0.181 ± 0.659 | 0.783 |
| log(Crop Size) | 0.277 ± 0.068 | **<0.001** |
| log(Pulp mass) | 0.486 ± 0.192 | **0.011** |
| log(Plant Area) | 0.959 ± 0.099 | **<0.001** |
| Site | 0.751 ± 0.134 | **<0.001** |

### H. Null models for interaction asymmetry estimates

In order to determine if matrix size was having an effect on the asymmetry distribution values encountered, we repeated the analysis subsampling from the total number of plants. Asymmetry values could be affected by the number of plants selected and sampled in the study because of varying matrix size and shape. We considered three different matrix sizes, of 20, 40 and 60 plants, that were compared to the asymmetry obtained from the 80 plants observed matrix. We performed 1000 permutations for each matrix dimension. Asymmetry in subsampled matrices was not greatly altered (Fig. S.H.1). All matrices showed few symmetric interactions. However, when the matrix included fewer plants the frequency of interactions where the animal is more dependent on the plant (i.e. negative asymmetries towards -1) increased while the frequency of interactions where the plant is more dependent on the animal (i.e. positive asymmetry values towards +1) decreased. This change in the sign of asymmetry is expected, given that a reduction in the number of plants available would lead to a greater estimated dependence of the birds on individual plants.

In addition, to test whether the asymmetry distribution encountered deviates from the expected asymmetry in randomly-built matrices, we compared the observed values to those obtained with null model matrices. We randomised fruit consumption following both Patefield and Vázquez null models (n=1000 permutation per model) (Dormann *et al.* 2009). The asymmetry frequency distribution encountered with both null models also maintained a "U" shaped pattern (Fig. S.H. 2). The Patefield null model increased the total number of unique-pairwise interactions, as it does not constrain connectance, allowing plants to create new links with birds and increasing their interaction degree. The creation of new links also caused an increase in the cases where animals were more dependent on plants. On the contrary, Vázquez null model results did not differ from the observed asymmetry distribution, except on a slight but significantly lower number of interactions for the more dependent avian species. That is, our system presented a higher frequency of interactions in which the animal is more dependent than would be expected when maintaining network connectance and species were allowed to interact randomly.

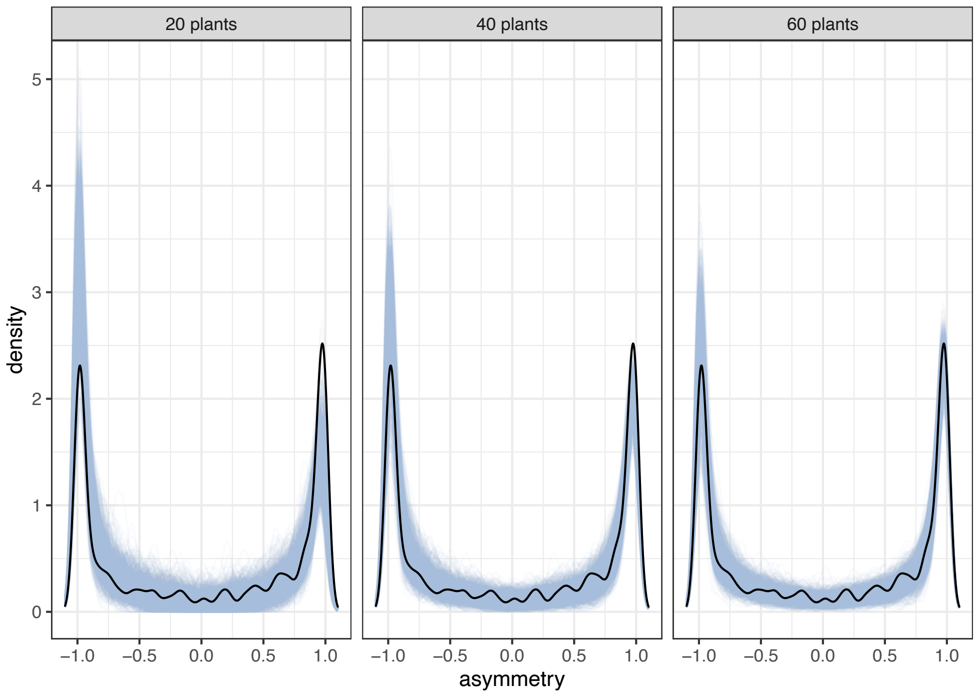

**Figure S.H.1.** Frequency distributions (density function) for interaction asymmetry when using three different matrix sizes (i.e. reducing rows to 20, 40 or 60 individual plants). Thin blue lines represent the 1000 permutations per matrix dimension. Black line represents the median observed asymmetry values.

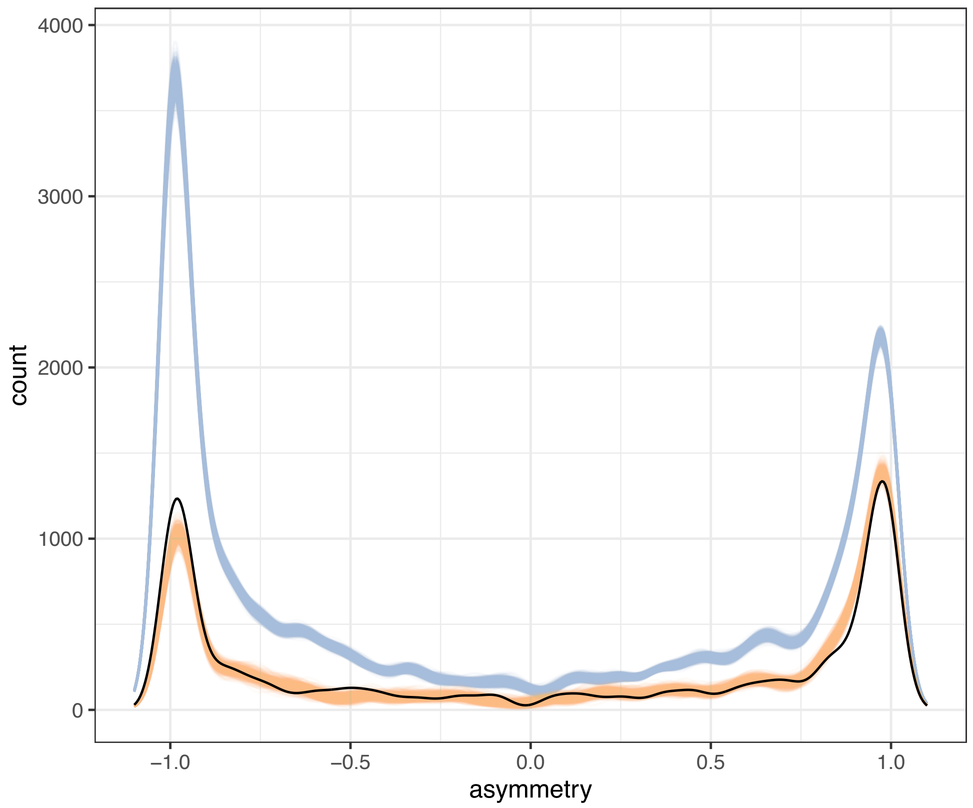

**Figure S.H.2.** Frequency distribution (count data) for interaction asymmetry using counts for null models. The blue line represents the 1000 permutations of the Patefield model; the orange line represents the 1000 permutations of the Vázquez model; black line represents median observed asymmetry values.

### I. Software

We used R version 4.1.2 (R Core Team 2021) and the following R packages: bayesplot v. 1.8.1 (Gabry et al. 2019; Gabry and Mahr 2021), bayestestR v. 0.11.5 (Makowski, Ben-Shachar, and Lüdecke 2019), bipartite v. 2.16 (Dormann, Gruber, and Fruend 2008), brms v. 2.16.3 (Bürkner 2017, 2018), DHARMa v. 0.4.5 (Hartig 2022), effect.lndscp v. 0.2.8 (Jordano and Rodriguez-Sanchez 2019), ggdist v. 3.1.1 (Kay 2022), ggpubr v. 0.4.0 (Kassambara 2020), ggridges v. 0.5.3 (Wilke 2021), glmmTMB v. 1.1.2.3 (Brooks et al. 2017), here v. 1.0.1 (Müller 2020), knitr v. 1.37 (Xie 2014, 2015, 2021), lme4 v. 1.1.28 (Bates et al. 2015), modelbased v. 0.7.2 (Makowski et al. 2020), patchwork v. 1.1.1 (Pedersen 2020), plotly v. 4.10.0 (Sievert 2020), rmarkdown v. 2.12 (Xie, Allaire, and Grolemund 2018; Xie, Dervieux, and Riederer 2020; Allaire et al. 2022), summarytools v. 1.0.0 (Comtois 2021), tidylog v. 1.0.2 (Elbers 2020), tidyverse v. 1.3.1 (Wickham et al. 2019), vegan (Oksanen et al. 2020), and visreg v. 2.7.0 (Breheny and Burchett 2017).

*R package citations*

Allaire, JJ, Yihui Xie, Jonathan McPherson, Javier Luraschi, Kevin Ushey, Aron Atkins, Hadley Wickham, Joe Cheng, Winston Chang, and Richard Iannone. 2022. *Rmarkdown: Dynamic Documents for R*. https://github.com/rstudio/rmarkdown.

Bates, Douglas, Martin Mächler, Ben Bolker, and Steve Walker. 2015. "Fitting Linear Mixed-Effects Models Using lme4." *Journal of Statistical Software* 67 (1): 1–48. https://doi.org/10.18637/jss.v067.i01.

Breheny, Patrick, and Woodrow Burchett. 2017. "Visualization of Regression Models Using Visreg." *The R Journal* 9 (2): 56–71.

Brooks, Mollie E., Kasper Kristensen, Koen J. van Benthem, Arni Magnusson, Casper W. Berg, Anders Nielsen, Hans J. Skaug, Martin Maechler, and Benjamin M. Bolker. 2017. "glmmTMB Balances Speed and Flexibility Among Packages for Zero-Inflated Generalized Linear Mixed Modeling." *The R Journal* 9 (2): 378–400. https://journal.r-project.org/archive/2017/RJ-2017-066/index.html.

Bürkner, Paul-Christian. 2017. "brms: An R Package for Bayesian Multilevel Models Using Stan." *Journal of Statistical Software* 80 (1): 1–28. https://doi.org/10.18637/jss.v080.i01.

———. 2018. "Advanced Bayesian Multilevel Modeling with the R Package brms." *The R Journal* 10 (1): 395–411. https://doi.org/10.32614/RJ-2018-017.

Comtois, Dominic. 2021. *Summarytools: Tools to Quickly and Neatly Summarize Data*. https://github.com/dcomtois/summarytools.

Dormann, C. F., B. Gruber, and J. Fruend. 2008. "Introducing the Bipartite Package: Analysing Ecological Networks." *R News* 8 (2): 8–11.

Elbers, Benjamin. 2020. *Tidylog: Logging for 'Dplyr' and 'Tidyr' Functions*. https://github.com/

elbersb/tidylog/.

Gabry, Jonah, and Tristan Mahr. 2021. "Bayesplot: Plotting for Bayesian Models." https://mc-stan.org/bayesplot/.

Gabry, Jonah, Daniel Simpson, Aki Vehtari, Michael Betancourt, and Andrew Gelman. 2019. "Visualization in Bayesian Workflow." *J. R. Stat. Soc. A* 182: 389–402. https://doi.org/10.1111/rssa.12378.

Hartig, Florian. 2022. *DHARMa: Residual Diagnostics for Hierarchical (Multi-Level / Mixed) Regression Models*. http://florianhartig.github.io/DHARMa/.

Jordano, Pedro, and Francisco Rodriguez-Sanchez. 2019. *Effect.Indscp: Effectiveness Landscapes*. https://github.com/pedroj/effectiveness_pckg

Kassambara, Alboukadel. 2020. *Ggpubr: 'Ggplot2' Based Publication Ready Plots*. https://rpkgs.datanovia.com/ggpubr/.

Kay, Matthew. 2022. *ggdist: Visualizations of Distributions and Uncertainty*. https://doi.org/10.5281/zenodo.3879620.

Makowski, Dominique, Mattan S. Ben-Shachar, and Daniel Lüdecke. 2019. "bayestestR: Describing Effects and Their Uncertainty, Existence and Significance Within the Bayesian Framework." *Journal of Open Source Software* 4 (40): 1541. https://doi.org/10.21105/joss.01541.

Makowski, Dominique, Mattan S. Ben-Shachar, Indrajeet Patil, and Daniel Lüdecke. 2020. "Estimation of Model-Based Predictions, Contrasts and Means." *CRAN*. https://github.com/easystats/modelbased.

Müller, Kirill. 2020. *Here: A Simpler Way to Find Your Files*. https://here.r-lib.org/

Oksanen, J., Blanchet, F.G., Friendly, M., Kindt, R., Legendre, P., McGlinn, D., et al. (2020). *vegan: Community Ecology Package*. R. https://CRAN.R-project.org/package=vegan

Pedersen, Thomas Lin. 2020. *Patchwork: The Composer of Plots*. https://patchwork.data-imaginist.com

———. 2021. *Ggforce: Accelerating 'Ggplot2'*. https://ggforce.data-imaginist.com

R Core Team. 2021. *R: A Language and Environment for Statistical Computing*. Vienna, Austria: R Foundation for Statistical Computing. https://www.R-project.org/.

Sievert, Carson. 2020. *Interactive Web-Based Data Visualization with R, Plotly, and Shiny*. Chapman; Hall/CRC. https://plotly-r.com.

Wickham, Hadley, Mara Averick, Jennifer Bryan, Winston Chang, Lucy D'Agostino McGowan, Romain François, Garrett Grolemund, et al. 2019. "Welcome to the tidyverse." *Journal of Open Source Software* 4 (43): 1686. https://doi.org/10.21105/joss.01686.

Wilke, Claus O. 2021. *Ggridges: Ridgeline Plots in 'Ggplot2'*. https://wilkelab.org/ggridges/.

Xie, Yihui. 2014. "Knitr: A Comprehensive Tool for Reproducible Research in R." In

*Implementing Reproducible Computational Research*, edited by Victoria Stodden, Friedrich Leisch, and Roger D. Peng. Chapman; Hall/CRC. http://www.crcpress.com/product/isbn/9781466561595.

———. 2015. *Dynamic Documents with R and Knitr*. 2nd ed. Boca Raton, Florida: Chapman; Hall/CRC. https://yihui.org/knitr/.

———. 2021. *Knitr: A General-Purpose Package for Dynamic Report Generation in r*. https://yihui.org/knitr/.

Xie, Yihui, J. J. Allaire, and Garrett Grolemund. 2018. *R Markdown: The Definitive Guide*. Boca Raton, Florida: Chapman; Hall/CRC. https://bookdown.org/yihui/rmarkdown.

Xie, Yihui, Christophe Dervieux, and Emily Riederer. 2020. *R Markdown Cookbook*. Boca Raton, Florida: Chapman; Hall/CRC. https://bookdown.org/yihui/rmarkdown-cookbook.
